# Cytological and preliminary genomic analysis of two *Leptodactylus* frog species (Anura, Leptodactylidae) with recently evolved large meiotic rings of multiple X and Y sex chromosomes

**DOI:** 10.1101/2025.03.27.645801

**Authors:** Jhon Alex Dziechciarz Vidal, Deborah Charlesworth, Wen-Juan Ma, Qi Zhou, Ricardo Utsunomia, Anderson José Baia Gomes, Amanda Bueno da Silva, Fábio Porto-Foresti, Thomas Liehr, Marcelo de Bello Cioffi

**Affiliations:** Laboratory of Evolutionary Cytogenetics, Department of Genetics and Evolution, Federal University of São Carlos, São Carlos, SP, Brazil; Institute of Ecology and Evolution, Ashworth Laboratories, King’s Buildings, University of Edinburgh, West Mains Road, EH9 3FL, Edinburgh, UK; Research group of Ecology, Evolution and Genetics, Biology Department, Vrije Universiteit Brussel, Brussels, Belgium; Life Sciences Institute, Zhejiang University, Hangzhou, Zhejiang, China; Faculdade de Ciências, UNESP, Bauru, São Paulo, Brazil; Laboratório de Biologia Molecular, Evolução e Microbiologia, Instituto Federal do Pará, Abaetetuba, Brazil; Jena University Hospital, Friedrich Schiller University, Institute of Human Genetics, Jena, Germany

**Keywords:** Meiotic multivalents, Repetitive DNAs, Translocations, Satellitome, Leptodactylidae

## Abstract

A few species have evolved multiple sex chromosome systems with more than two Xs or Ys. These involve sex chromosome-autosome translocations (sometimes called fusions as very small heterochromatic arms may be deleted), creating neo-sex chromosome systems. Among vertebrates, frogs (Anura) have the highest known number of such translocation systems. This study within the genus *Leptodactylus*, investigated the two species *L. pentadactylus* (LPE) and *L. paraensis* (LPA), in which large ring multivalents are seen in male meiosis, indicating translocations involving the sex chromosomes. Four other species studied do not have such rings, but they share characteristics making rearrangements less likely to be eliminated. To start understanding the formation of multivalents, we used genomic and cytogenetic methods to investigate repetitive DNA sequences, including satellite DNAs, rDNAs, and telomeric sequences, and conducted comparative genomic hybridization (CGH). The LPE genome includes a large number of satDNA families, and in situ mapping of several satDNAs individually identified eight of the ten chromosomes in its multivalent. In LPA, morphological similarities indicate that several chromosomes are shared by the multivalents of both species, and a candidate ancestral sex chromosome pair could be identified. *In situ* mapping in LPE suggests recent satDNA accumulation in the subtelomeric regions, which differ from those in the outgroup species, contrary to the expectation that the translocations create sex-linkage in the pericentromeric regions.

## Introduction

Sex chromosomes have evolved independently in many groups of organisms, and the degrees of differentiation between members of the pairs vary greatly across species, reviewed in ^1^. In animals, most species have XX/XY or ZZ/ZW sex chromosome systems (male and female heterogamety, respectively, and sometimes the Y or W lacks (almost) all functional genes, termed “highly degenerated”). A minority have multiple sex chromosome systems reflecting translocations between autosomes and one or both members of a pair of ancestral sex chromosomes, or fusions, which involve translocations when one arm is small and heterochromatic^2–4^. Reciprocal translocations are expected to greatly reduce fertility in heterozygotes^2^ and many translocations will thus fail to spread in populations unless selection favours the new arrangement^5^. Nevertheless, many systems with more than one X or Y chromosome, include rodents^6–9^, monkeys^6^, fish (reviewed in ^4^) and plants^7,8^; same can be seen for more than one Z chromosome in Lepidoptera, with ZZ/ZW systems^9–11^. Many systems involve a single pair of autosomes (♀X1X1X2X2/♂X1X2Y, ♀XX/♂XY1Y2, ♀X1X1X2X2/♂X1Y1X2Y2), or, perhaps less often, similar situations can be observed with female heterogamety^4,12,13^.

A small handful of species have evolved systems involving more than one autosome as well as the sex chromosome pair. All known multiple systems with meiotic chains or rings are derived from XY systems^14–17^, including two Monotreme (non-Eutherian) mammals, the platypus *Ornithorhynchus anatinus* and the short-beaked echidna *Tachyglossus aculeatus*, both with X1-X5/Y1-Y5 systems with multivalent chains of 10 chromosomes in male meiosis^14,18,19^. The evolution of these systems is not yet well understood, though it is clear that the components that show Y linkage are fully Y-linked, as they have become genetically degenerated in both species and have lost most genes carried on their X counterparts^20^. Studying a younger system may provide information about the time course of degeneration and possibly the reason for these systems’ formation.

Among other vertebrate clades, frogs include species with chain or ring multivalents. Three cases have currently been described. Males of the smoky jungle frog *Leptodactylus pentadactylus* (hereafter denoted by LPE) have the largest multiple sex chromosome system so far identified in any vertebrate, a ring of 12 chromosomes that are largely unpaired in male meiosis (an X1-X6/Y1-Y6 system, see in ^15,17^, though some populations of this species have a ring of 10^21^. Large meiotic rings have been described in two other frogs distantly related to *Leptodactylus*: the clay robber *Haddadus binotatus* (Craugastoridae)^22^ and Taiwanese frog *Odorrana swinhoana*^23^, with rings of 8 and 6 chromosomes, respectively. The two *Leptodactylus* species studied here, LPE and *L. paraensis* (abbreviated to LPA), separated about 28 million years ago (MYA), and diverged from the other two frog species approximately 86 and 150 MYA, respectively^24^; thus, meiotic rings most likely emerged independently in these three frog lineages.

As so few species have multi-chromosomal chains or rings with more than four sex chromosomes (reviewed in ^14^), frogs might have an unusual tendency to undergo multiple rearrangements involving the sex chromosomes, perhaps because their chromosomes have suitable properties (see below). Frogs might thus be suitable for studying the rates of appearance of X- and/or Y-autosomal translocations and meiotic rings, and, if they show an increased incidence compared with other organisms, what factors lead to this. In the case of rings involving the sex chromosomes, translocations might be an evolutionary response to a situation in which recombination is disfavoured. If crossovers are strictly localized to chromosome tips and segregation of adjacent centromeres in meiosis is alternating, a translocation can cause the middle chromosome regions to become fully sex-linked. An autosomal sexually antagonistic polymorphism might thereby become fully sex-linked via mutual translocation with part of the sex-linked region^25^. However, testing the hypothesis that such selection favours an observed lack of recombination between the sex chromosomes is notoriously difficult.

Before considering factors that might favour reciprocal translocations, it is important to consider the genomic and ecological contexts that affect the probability of such rearrangements establishing. Genomic factors might include characteristics that (i) increase the probability of a reciprocal translocation arising or (ii) might decrease the deleterious consequences of chromosomal breaks (such as frequent unbalanced gametes or disrupting the effects of upstream regulatory sequences on the expression of genes) so that a translocation is more likely to be able to increase in frequency in a population. The two meiotic characteristics mentioned above are important in this category. Also, (iii) ecological characteristics, such as particularly small and isolated populations, can create a higher chance of genetic drift fixing the translocations.

The category (i) has been the subject of numerous studies testing for fragile regions, particularly prone to chromosomal breakage, in the genomes of certain eukaryotic species. These regions are often referred to as “evolutionary breakpoint regions” (EBRs) to denote their tendency to experience chromosome breakage in independent lineages. Analysis of numerous eukaryotic genomes indicates that EBRs and fragile sites in specific species tend to be regions with particularly high repeat richness, such as pericentromeric and subtelomeric regions, rather than specific genomic sites (reviewed in ^26^). This might be expected, since repetitive sequences can produce DNA structures, such as hairpins, which may facilitate chromosomal rearrangements and can recombine “ectopically” with similar sequences in other genome regions^27,28^. On the other hand, open chromatin, which makes breaks more likely, is associated with high densities of active genes and high recombination rates. Such genomic properties can account for convergent breakpoints in independent lineages. In the case of species with reciprocal translocations creating rings, the breakpoints are most likely to be in highly repetitive middle regions of metacentric chromosomes in the ancestors. However, regions with highly repetitive sequences may appear in new genomic locations, including neo-centromeres (as reviewed in ^29^), or be lost from existing locations, causing turnovers of repeat content, especially in rarely recombining regions^30^, so different rearrangement breakpoint sites are possible in related species^27^.

Category (ii) includes genomic factors that, rather than increasing the probability that rearrangements arise, decrease their deleterious consequences. This category includes weak purifying selection, leading to EBRs in gene-dense, recombining parts of genomes being mainly in permanent genomic features such as intergenic regions (as reviewed in ^26^). Category (ii) also includes factors reducing the frequency of unbalanced gametes, such as localization of crossovers mainly to chromosome tips and alternating segregation of adjacent centromeres in the multivalent in meiosis. Importantly, if the middle chromosome regions that rarely cross over are repeat-rich in the ancestral species, this characteristic will usually be inherited by the rings. Conversely, repeat-richness (a category (i) characteristic causing a high origination rate of rearrangements) correlates with regions exhibiting low recombination rates and low gene densities, as the absence of recombination facilitates ectopic exchanges, enabling rearrangements to occur without inducing meiotic adverse effects.

Category (iii) includes characteristics affecting the probability that individuals in a population will be heterozygous for a translocation. A newly arisen translocation will initially be almost exclusively heterozygous, and such heterozygotes often suffer the well-known disadvantages of unbalanced gamete production (reviewed in ^2,31^). In small and isolated populations, however, genetic drift may allow such a rearrangement to reach a high enough frequency to avoid being lost and sometimes to fix^32^. Inbreeding similarly allows translocations to spread in a population, simply because heterozygotes are rare, and indeed, there is evidence for higher chromosome rearrangement rates in inbreeders (reviewed in ^33^).

Out of the ∼120 *Leptodactylus* species, 45 have been analysed cytogenetically. Most species have 2n = 22 chromosomes. As only a few studies have examined meiosis (all conducted in males), it is unclear how many species might also have systems with chromosomal chains or rings. A previous study showed that, in LPE, segregation of adjacent centromeres alternates in male meiosis, a category (ii) factor that facilitates the production of balanced gametes^34^. We return to the other categories in the Discussion section, but here we are concerned mainly with studying genomic category (i) factors in the genus *Leptodactylus* frogs and the build-up of the ring system, using combined genome sequencing and cytogenetic *in situ* hybridization analysis of several species. A goal was to describe repetitive DNA sequences (largely satellite sequences, plus some preliminary results from transposable elements) and evaluate changes during the evolution of the large chromosomal ring multivalents, and to ask whether EBRs might be involved, using two species that both possess rings.

A phylogenetic tree of five species in the genus is shown in **Supplementary** Figure 1. Our study confirmed the large ring systems and different autosome numbers already recognized in LPE. Among the six species examined cytologically, only two have ring sex chromosomes, LPE and LPA, whose chromosomes have not been studied previously, but which proved to have multiple sex chromosomes (**Table 1, Supplementary Table 1)**. The phylogeny suggests that the ring sex chromosome of LPE evolved after the split from *L. labyrinthicus,* which does not have ring chromosomes. This split is estimated to have occurred about 10 million generations ago, as the species is thought to have one generation per year^35^. The current phylogeny excludes LPA, for which sequence data are unavailable. Another objective of this study was to determine whether the translocations evolved independently in the LPE and LPA lineages or whether some evolved in a common ancestor. Our study included a new LPE population, which revealed a different number of chromosomes in the ring, showing that new translocations are still possible. This is also consistent with evidence that LPE is a species complex^15,36^; hybrids may form in contact zones between its members. As a first step towards understanding whether the LPE and LPA species’ rings evolved independently, and to help identify their breakpoints, we identified most of the chromosomes in the LPE ring.

**Table 1:**
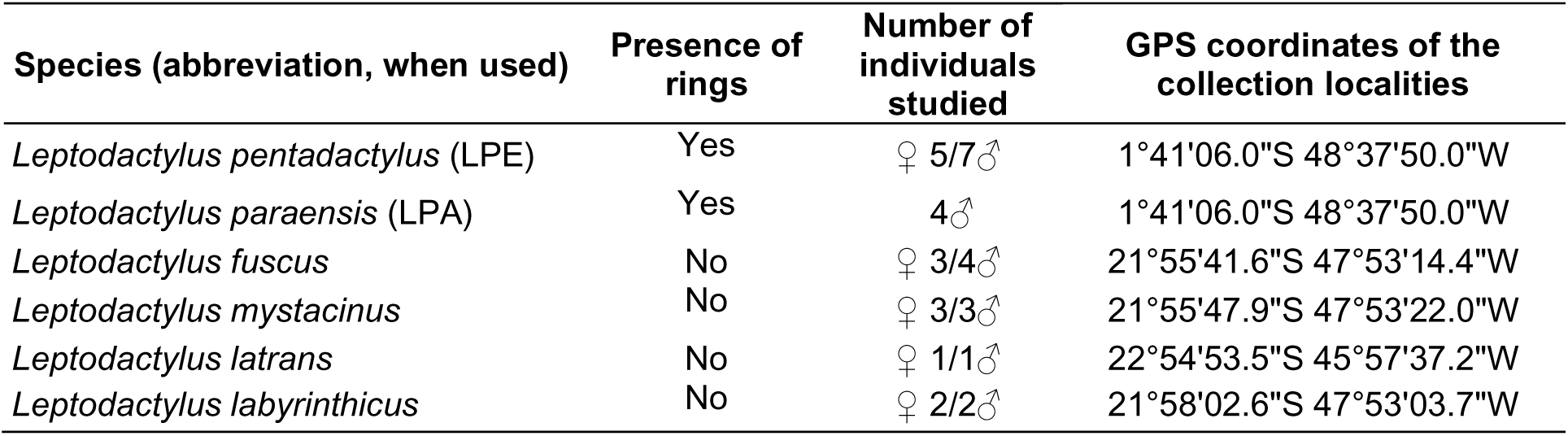
*Leptodactylus* species studied here, including the numbers of individuals and geographic coordinates of the sampling sites.

A final goal was to ask whether the translocations caused the autosomal regions added to the rings to become completely sex linked. This can potentially be tested by asking whether repetitive sequence content has increased in the autosomal regions that were translocated and became members of the rings. However, as mentioned above, the non-terminal regions of these chromosomes may have already been rarely recombining in the ancestor. It may therefore be impossible to distinguish between high repetitive sequence content at the base of chromosome arms constituting a category (i) factor creating a high rearrangement rate, versus the translocation events that caused sex-linkage having reduced these regions’ recombination rates, or increased the size of the regions in which recombination rates are low.

## Results

### Cytogenetic analyses

We examined mitosis and meiosis in LPE. Both males and females displayed the expected metacentric or sub-metacentric chromosomes (2n=22, see (**Figure 1a, c**). All five males investigated showed heteromorphism for chromosomes 3 and 5. No heteromorphism was detected in any of the five females analysed for these or any other chromosome pair **(Figure 1c, d)**. Specifically, the chromosome 3 and 5 short arms are longer than that of the non-male-specific homologue, as expected if these arms have stopped recombining and started accumulating repeats **(Figure 1a, b)**. This observation (and our satDNA mapping—see below) suggests that these two chromosomes are part of the multiple LPE sex chromosome system. Differentiation of this heteromorphic chromosome pair suggests that recombination has been suppressed for long enough to allow considerable accumulation of repetitive sequences (resulting in length differences from the homologs), though neither is heterochromatic. One or the other of these heteromorphic chromosomes may represent the ancestral XY pair, but our current analyses do not allow us to determine which one.

**Figure 1.**
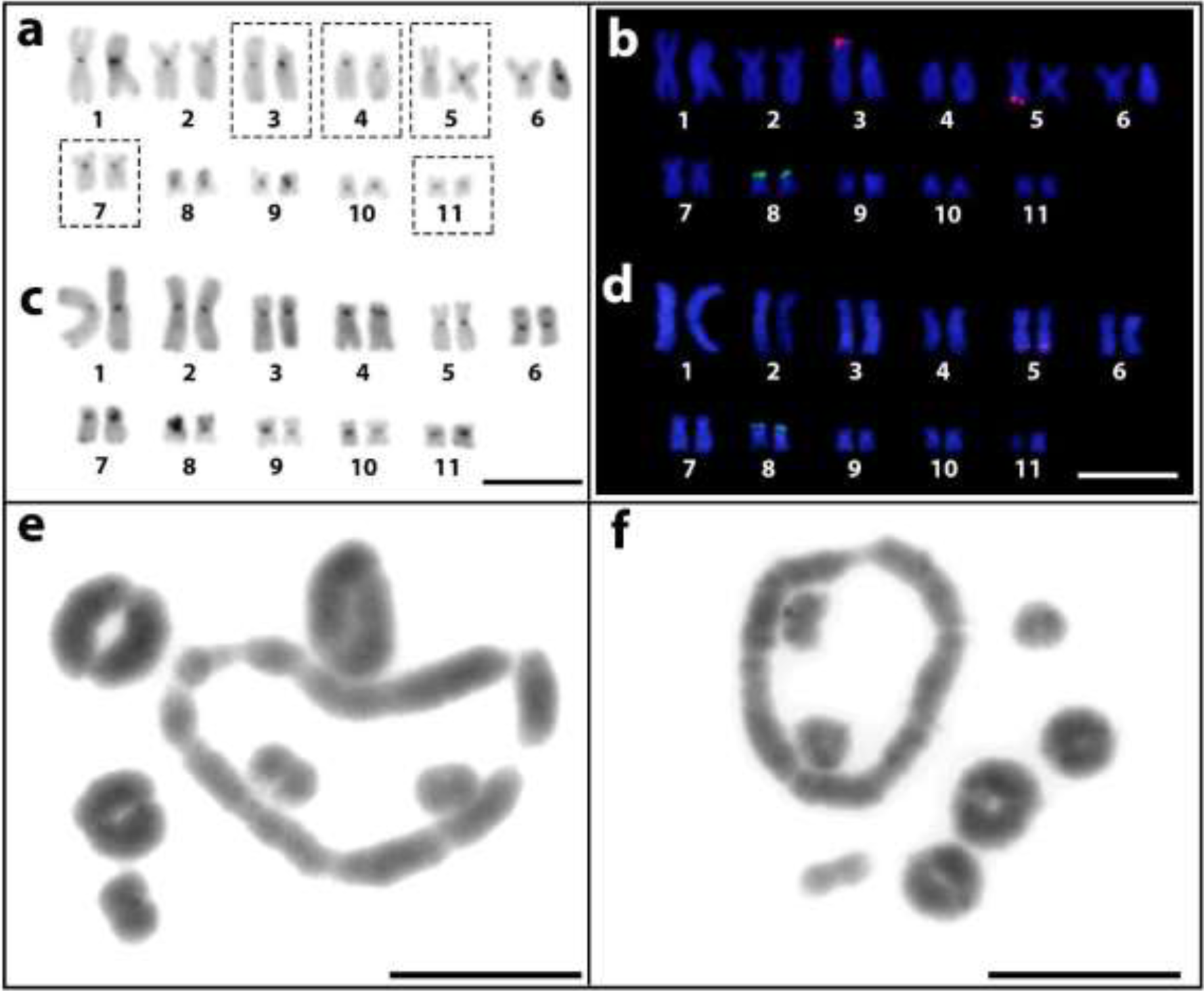
Karyotypes of male **(a and b)** and female **(c and d)** of LPE analysed by C-banding to show heterochromatic regions, including centromeric regions in the middle of each chromosome **(a and c)** and FISH with two probes, 18S and 5S rDNA, in green and red, respectively **(b and d)**. Metaphase I shows six bivalents and a ring of 10 chromosomes in LPE, and a ring of eight chromosomes and seven bivalents in LPA, respectively (**e and f)**. The dotted frames indicate the chromosomal pairs inferred to be involved in the rings based on our *in situ* mapping results described below. Bars = 10um.

Males of LPA were examined in meiosis (metaphase I and II). As in LPE, multi-chromosomal rings were seen, and both the rings and univalents segregated in meiosis II, producing products with the expected n=11 (**Supplementary** Figure 3). LPE has a ring of 10 chromosomes (including about 42% of the genome, based on the lengths in **Figures 1a and b**) plus six bivalents, while LPA had a ring of 8 (about 30% of the genome, based on **Figure 1f**), and seven bivalents **(Figures 1e and f)**. In both species, all bivalents (not only the multivalents) form rings during male meiosis **(Figures 1e and f)**, suggesting that pairing, and probably crossing over, are strongly localized to the tips of all chromosomes in LPE and LPA karyotypes.

We analysed mitotic and male meiotic chromosomes in the other four *Leptodactylus* species **(Table 1).** All showed 2n = 22, corroborating previous descriptions for this species^17,37,38^. In meiosis, none of these four species have multivalents. Instead, eleven bivalents are seen, which formed rings during male meiosis, supporting the conclusion just mentioned that crossover localization to chromosome tips is ancestral in this genus **(Supplementary** Figure 4). C-positive heterochromatin was previously detected in the centromeric regions of all chromosomes of all these species^17^, and additional heterochromatic blocks were seen in the terminal regions of LPE pairs 6, 7, and 8 in both sexes **(Figure 1a and c).**

### Repetitive DNA composition of L. pentadactylus *and* L. fuscus and characterization of LpeSatDNAs

The *de novo* TE annotation RepeatModeler2 pipeline estimated that nearly 58.5% of the *L. fuscus* genome is composed of TEs, comparable to the available estimates for closely related frog lineages, including Leptodactylidae with about 63% and Hylidae with 65%^39^. The estimated overall repetitive DNA content (including TEs and other repeats) in LPE was lower, at approximately 43% and 42% in males and females, respectively, based on the *L. fuscus* library **(Supplementary** Figure 5). It was around 50% using DNApipeTE **(Supplementary** Figure 6), which may more accurately reflect the actual abundance, as it is designed for low-coverage Illumina short reads without genome assembly information. However, the estimated total amounts cannot reliably be compared because the LPE data were analyzed using a repeat library from the distantly related *L. fuscus*, although most of the main TE classes were identified in both species, with DNA transposons the most prevalent **(Supplementary** Figures 5 and 6). An analysis of the divergence landscape in *L. fuscus* suggests that TEs are currently active, though some insertions are older and established, a pattern that is frequently seen. For LPE, the landscape is nearly ‘L’ shaped, suggesting the possibility that a higher proportion of TE insertions, especially DNA transposon types, occurred more recently than in *L. fuscus* **(Supplementary** Figure 5). However, this type of analysis has little power to resolve ages of elements, especially given that the LPE data were analyzed using the *L. fuscu*s repeat library. Most repetitive sequences were categorized as unknown or non-annotated, potentially indicating species-specific sequences, but we cannot exclude the possibility that the low repetitive percentage may merely stem from insufficient coverage of the short reads. Overall, we have no direct evidence associating the multivalent formation with the expansion of TEs in LPE since its split from its sister species, *L. labyrinthicus* (which has no ring sex chromosomes).

We next examined satellite DNAs, which can sometimes cause expansions of genome regions that recombine rarely, particularly pericentromeric regions, as found in *Drosophila* species^29^. In the nearly 1x coverage representative LPE genome sequences (see Methods), we identified 104 satDNA sequences, hereafter termed LpeSatDNAs, many of them A+T rich (range 21% to 83% AT, average 56%, as is frequently found). Most LpeSatDNA repeat units were >100 bp; their lengths range from 21 bp to 6,930 bp (for LpeSat07-21 and LpeSat16-6930, respectively), and the average was 429 bp. All 104 LpeSatDNAs were found in both sexes, but in different abundances **(Supplementary** Figure 8). Eight of these sequences, LpeSat34-31, LpeSat63-50, LpeSat72-70, LpeSat80-22, LpeSat91-37, LpeSat96-32, LpeSat100-390, and LpeSat101-74, were enriched in the male DNA sequences (mostly low abundance families), and five with higher abundances (LpeSat03-152, LpeSat36-325, LpeSat46-108, LpeSat48-178, and LpeSat74-403) in the female one (Figure 2**; Supplementary Table 2)**.

**Figure 2.**
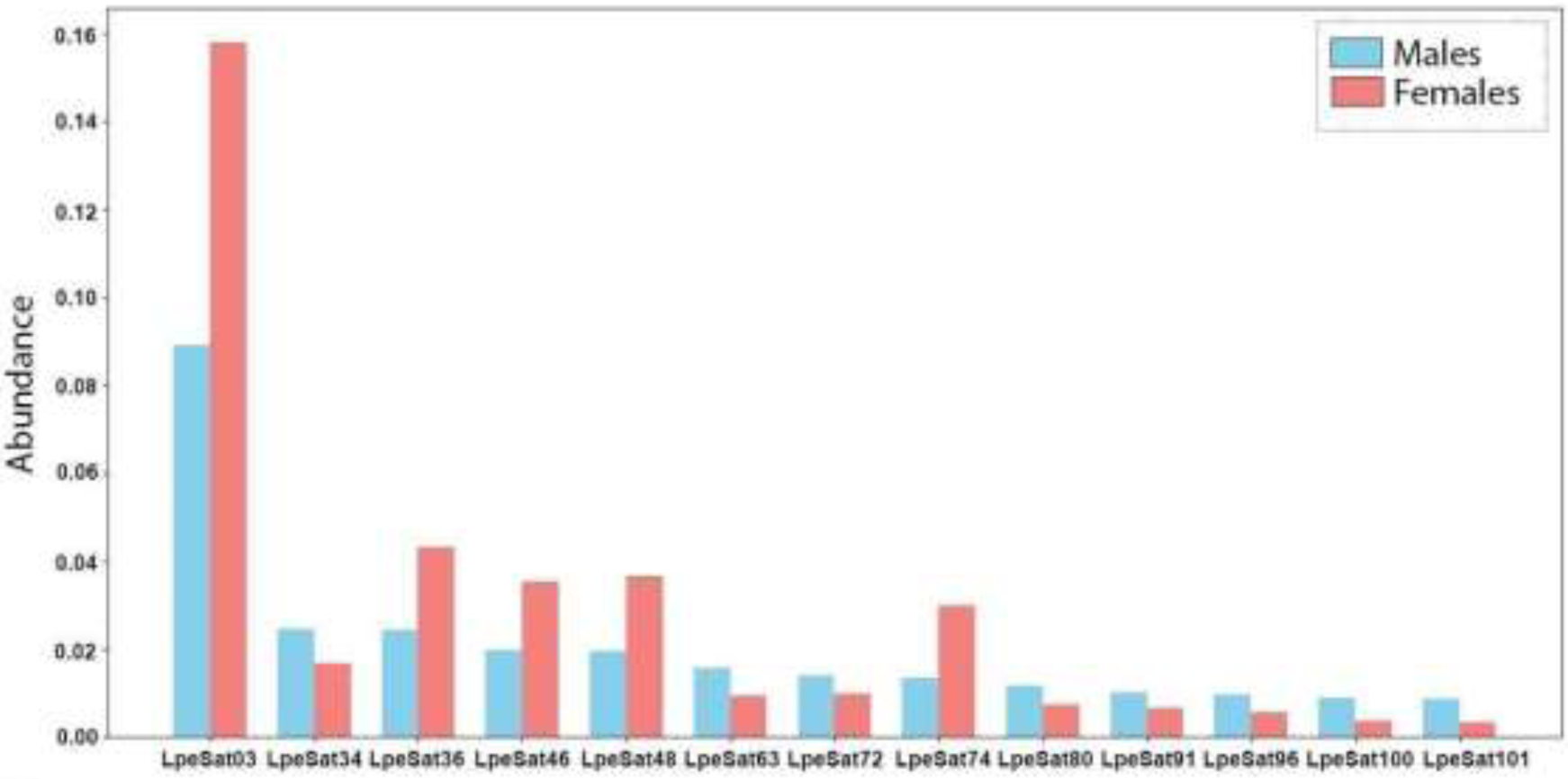
LpeSatDNAs with different abundances in males and females. The Y axis shows the estimated abundance of each family named on the X axis.

### In silico and in situ mapping of selected LpeSatDNAs against the L. fuscus genome

In the *L. fuscus* (GCA_031893025.1) genome assembly, mapping LpeSatDNA sequences detected only four out of the 104 LpeSatDNAs, indicating that these repeat sequences are ephemeral and turn over between species within the genus, with only some sequences being preserved over the estimated ∼28 MYA separating *L. pentadactylus* and *L. fuscus*. Among the four shared LpeSatDNAs, only LpeSat01-35 (by far the most abundant one) yielded positive hybridization signals in our *in situ* mapping experiments, mostly in or near the terminal regions of all chromosomes **(Supplementary** Figure 8).

### Mapping of LpeSatDNAs on LPE mitotic chromosomes

Eleven of the 18 LpeSatDNAs for which primers were designed (see Methods) showed positive hybridization signals on mitotic chromosomes of both LPE males and females (Figures 3 **and 4, Table 2**). LpeSat01-35 was again detected in terminal regions of all chromosomes, and LpeSat03-152 in centromere-proximal regions. LpeSat02-35 and LpeSat04-92 showed signals on almost all chromosomes at their terminal and centromere-proximal regions, respectively. We detected hybridization signals of LpeSat09-1589 in the pericentromeric regions of two chromosome pairs within the ring multivalent. LpeSat36-325, LpeSat48-178, LpeSat74-403, LpeSat91-37, and LpeSat100-390 hybridized with a single pair of chromosomes, while LpeSat80-22 was present in a single homolog from pair 3 (Figures 3 **and 4).**

**Figure 3.**
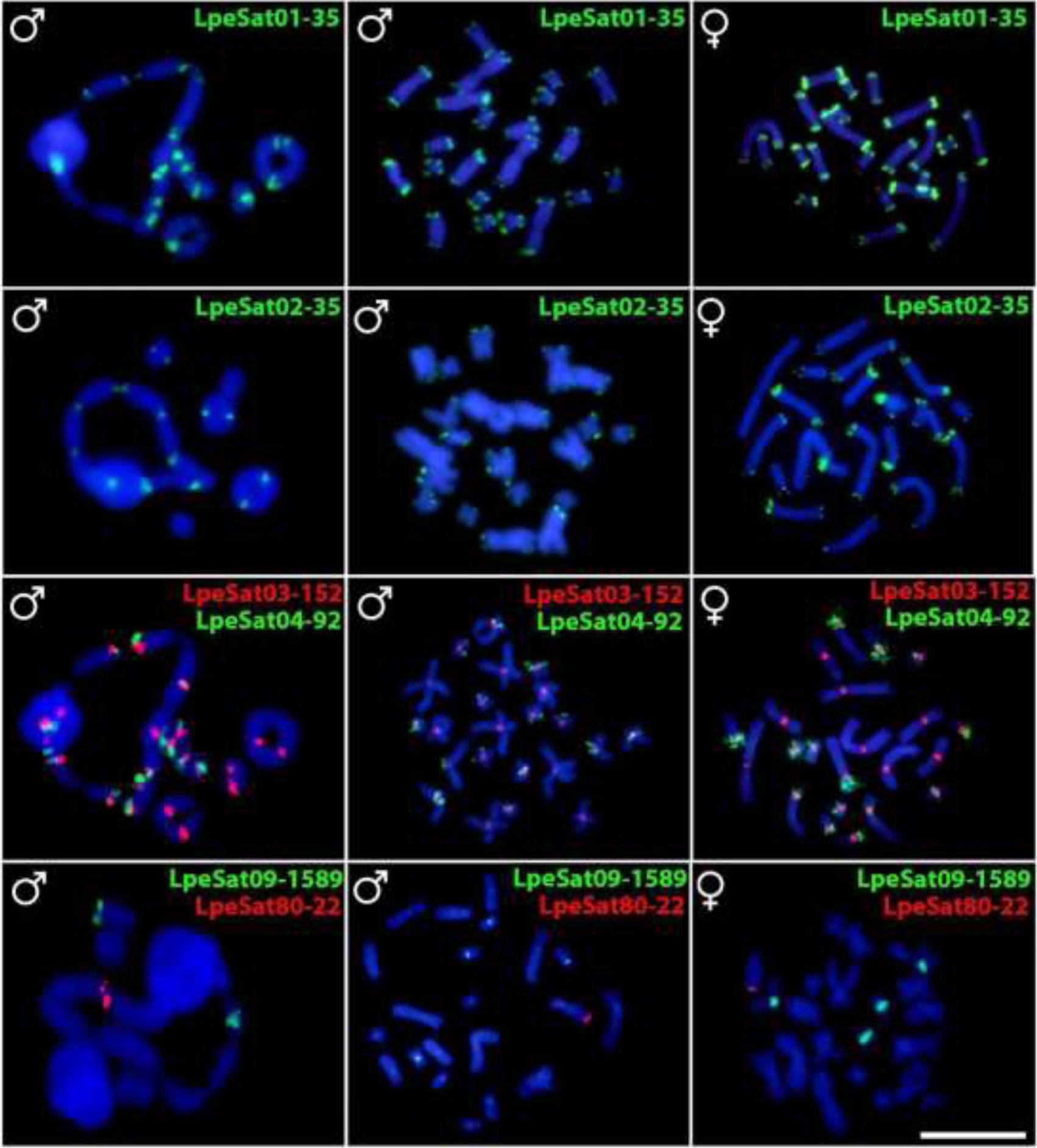
Male LPE meiotic cells (first column) and mitotic metaphases of a male and female (second and third columns, respectively) after FISH with the LpeSatDNA families indicated; these were detected on both autosomes and sex chromosomes. Green indicates staining with Atto-488-dUTP, and red indicates Atto-550-dUTP staining. Bar = 10 μm.

**Figure 4.**
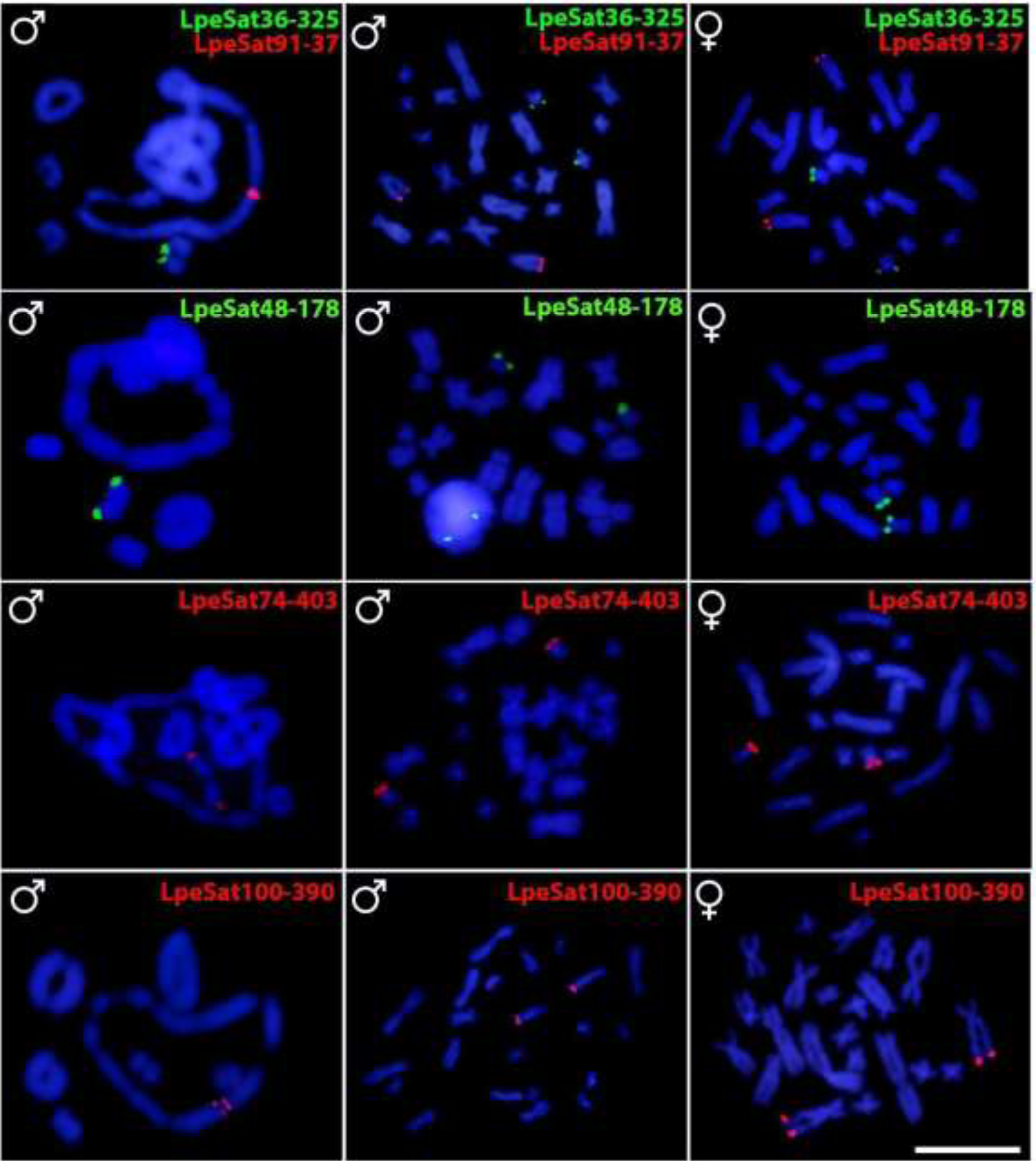
Male LPE meiotic cells (first column) and mitotic metaphases of a male and female (second and third columns, respectively) after FISH with the LpeSatDNA monomers indicated; these were also detected on both autosomes and sex chromosomes. Green indicates staining with Atto-488-dUTP, and red indicates Atto-550-dUTP staining. Bar = 10 μm.

**Table 2:** Chromosomal distribution of DNA repeats (satDNA and rDNA sequences) on the LPE and LPA chromosomes. The DNA repeats that allowed us to determine the identities of individual chromosomes in the LPE decavalent meiotic ring are in bold font. The regions where repeats were detected are indicated as follows: telomeric regions (t), centromeric regions (c), and pericentromeric regions (pc).

| Numbers of chromosomes with hybridization sites |  |  |
| --- | --- | --- |
| DNA repeat | Locations in LPE | Locations in LPA |
| LpeSat01-35 | All 10 chromosomes in the ring<br>+ 6 bivalents (t) | All 8 chromosomes in the ring<br>+ 7 bivalents (t) |
| LpeSat02-35 | All 10 chromosomes in the ring<br>+ 4 bivalents (t) | 4 chromosomes in the ring<br>+ 2 bivalent (t) |
| LpeSat03-152 | All 10 chromosomes in the ring<br>+ 6 bivalents (c) | 2 chromosomes in the ring<br>+ 1 bivalent (c) |
| LpeSat04-92 | All 10 chromosomes in the ring<br>+ 2 bivalents (pc and t) | All 8 chromosomes in the ring<br>+ 7 bivalents (pc and t) |
| **LpeSat09-1589** | 2 chromosomes in the ring (**pair 7**)<br>+ 2 bivalents (pc and t) | No signal |
| LpeSat36-325 | Only a single bivalent (t) | No signal |
| LpeSat48-178 | Only a single bivalent (t) | No signal |
| LpeSat74-403 | Only a single bivalent (t) | No signal |
| **LpeSat80-22** | Only 1 chromosome in the ring (one<br>homolog of chromosome **pair 3**) (t) | No signal |
| **LpeSat91-37** | Only 2 chromosomes in the ring<br>(**pair 4**) (t) | No signal |
| **LpeSat100-390** | Only 2 chromosomes in the ring<br>(**pair 3**) (t) | No signal |
| 18S rDNA | Only one bivalent (t) | Only one bivalent (t) |
| **5S rDNA** | Only 2 chromosomes in the ring<br>(single homologs of each of<br>chromosome **pairs 3 and 5**) (t) | Only one bivalent (t) |

### Mapping of rDNAs, telomeric repeats, and comparative genomic hybridization (CGH) on LPE mitotic chromosomes

The telomeric probe (see Methods) mapped to the terminal region of all chromosomes in both species, identifying the telomeres mentioned in what follows, and no interstitial telomeric sites were detected **(Supplementary** Figure 8). Only pair 8 of the LPE mitotic chromosomes carries 18S rDNA sites (Figure 1). Sequential hybridization using 18S rDNA, LpeSat09-1589, LpeSat36-325, LpeSat48-178, and LpeSat74-403 probes established that these sequences all occur on this chromosome **(Supplementary** Figure 10, **Table 2)**. In females, the 5S rDNA probe hybridized to both homologs of pair 5, while in males, it hybridized only in a single homolog of each of pairs 3 and 5 providing further evidence that one translocation involved these chromosomal pairs (Figure 1). Other than this, intraspecific CGH experiments in LPE to search for sex-specific sequences revealed only overlapping signals in the centromeric regions, indicating similarity between the male and female genomes **(Supplementary** Figure 11).

### Mapping of LpeSatDNAs on LPE and LPA meiotic chromosomes

FISH experiments revealed that 18S rDNA sites mapped to a single bivalent in meiotic preparations from both species, as does 5S rDNA in LPA, whereas, as just described, it is detected in the LPE ring of 10, on chromosome 5 in females, or 3 and 5 in males (Figure 5). We found three main patterns of LpeSatDNA hybridization: 1) Present both on bivalents and on one or more chromosomes in the ring (LpeSat01-35, LpeSat02-35, LpeSat03-152, LpeSat04-92, and LpeSat09-1589); 2) Present only on bivalents (LpeSat36-325, LpeSat48-178, and LpeSat74-403); and 3) Present exclusively on one or more chromosomes in the ring (LpeSat80-22, LpeSat91-37 and LpeSat100-390) (**Table 2**, Figures 3 **and 4**). Two of the four most abundant LpeSatDNAs showed hybridization signals on all the chromosomes (**Table 2**), butLpeSat02-35 mapped only to three bivalents and LpeSat004-92 to just one (Figure 5).

**Figure 5.**
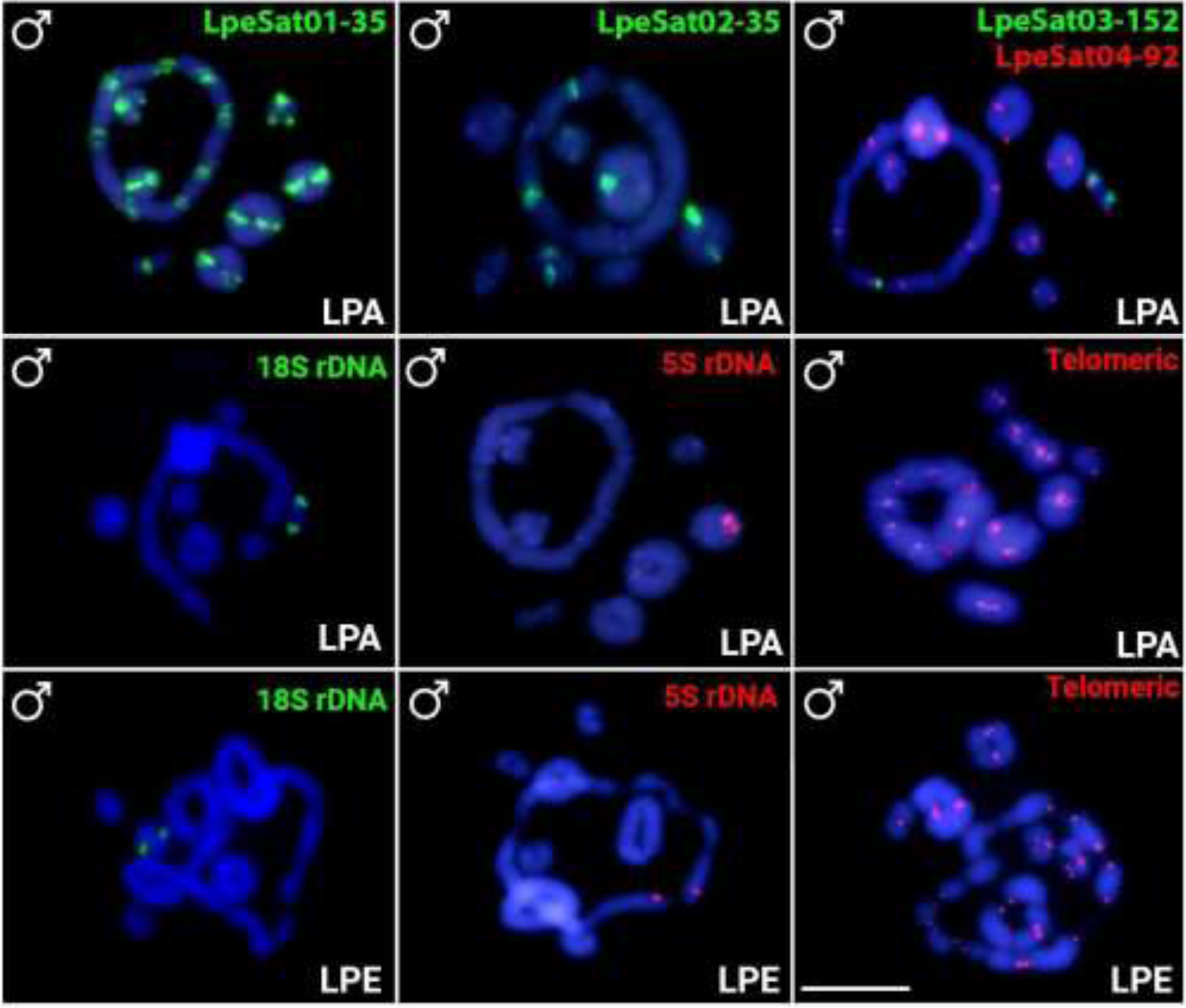
Male meiotic cells hybridized with LpeSatDNAs (LPA only) and rDNAS (both LPA and LPE). Green indicates staining with Atto-488-dUTP, and red indicates Atto-550- dUTP staining Scale bar = 10 μm.

### Identifying the chromosomes in the LPE meiotic ring

Despite using probe-specific FISH markers, the previous study^15^ was unable to identify the ancestral XY pair. Pair 3 was arbitrarily designated as X_1_ since it was the largest chromosome with no evident pair. Our *in situ* mapping of SatDNAs to meiotic and mitotic chromosomes of LPE allowed us to identify this pair, as well as pairs 4, 5, and 7, as four of the five chromosomes involved in the ring of 10 (Figure 1a, **Table 2).** Co-hybridization of LpeSat03-152 and LpeSat04-92 further showed that pairs 1 and 2 are not part of the decavalent, and hybridization in mitotic chromosomes shows that neither of these chromosomes exhibits co-localization of these satDNAs in their centromeric regions. Complementary meiotic investigations demonstrate that the two largest bivalents also lack this co-localization (Figure 3). The same applies to pair 8, a small bivalent that carries the 18S rDNA loci in this species (and also in LPA; Figure 5), confirming the previous inference in individuals with a ring of 12, whose chromosome pair carrying this locus was not in the multivalent^15,17^. Chromosomes 3, 4, and 7 are part of the ring of 10, as shown by the presence of LpeSat100-390 (on pair 3), LpeSat91-37 (pair 4), and LpeSat09-1589 (pair 7) within the multivalent (Figures 4 **and 5**). We also established that pair 5 is in the LPE multivalent, based on the presence of the 5S rDNA locus on this chromosome in the decavalent, and this is indicative of a rearrangement between pairs 3 and 5 (Figures 1 **and 5).** Based on its morphology, the final component of the ring-shaped multivalent was thought to be the smallest chromosome, pair 11.

None of the chromosomes within the multivalent could be definitively identified in the other ring-bearing species, LPA. However, their morphologies suggest that several chromosomes (e.g., pairs 3, 4 and 5) are shared by the LPE and LPA multivalents (Figures 1 **and 5).** As mentioned above, the 5S rDNA locus is not on a chromosome in the LPA ring, which suggests that pairs 3 and 5 may not represent the ancestral XY pair, despite their heteromorphism (Figure 1).

## Discussion and Conclusions

We confirm the presence of large ring multivalents in male meiosis of two *Leptodactylus* species, involving at least four X and Y-autosome reciprocal translocations. While LPE populations exhibiting rings of 12 chromosomes have been documented^15,17^, we now describe a population with only 10 chromosomes in the ring from a nearby location, similar that described by ^21^ (Figure 6).

**Figure 6.**
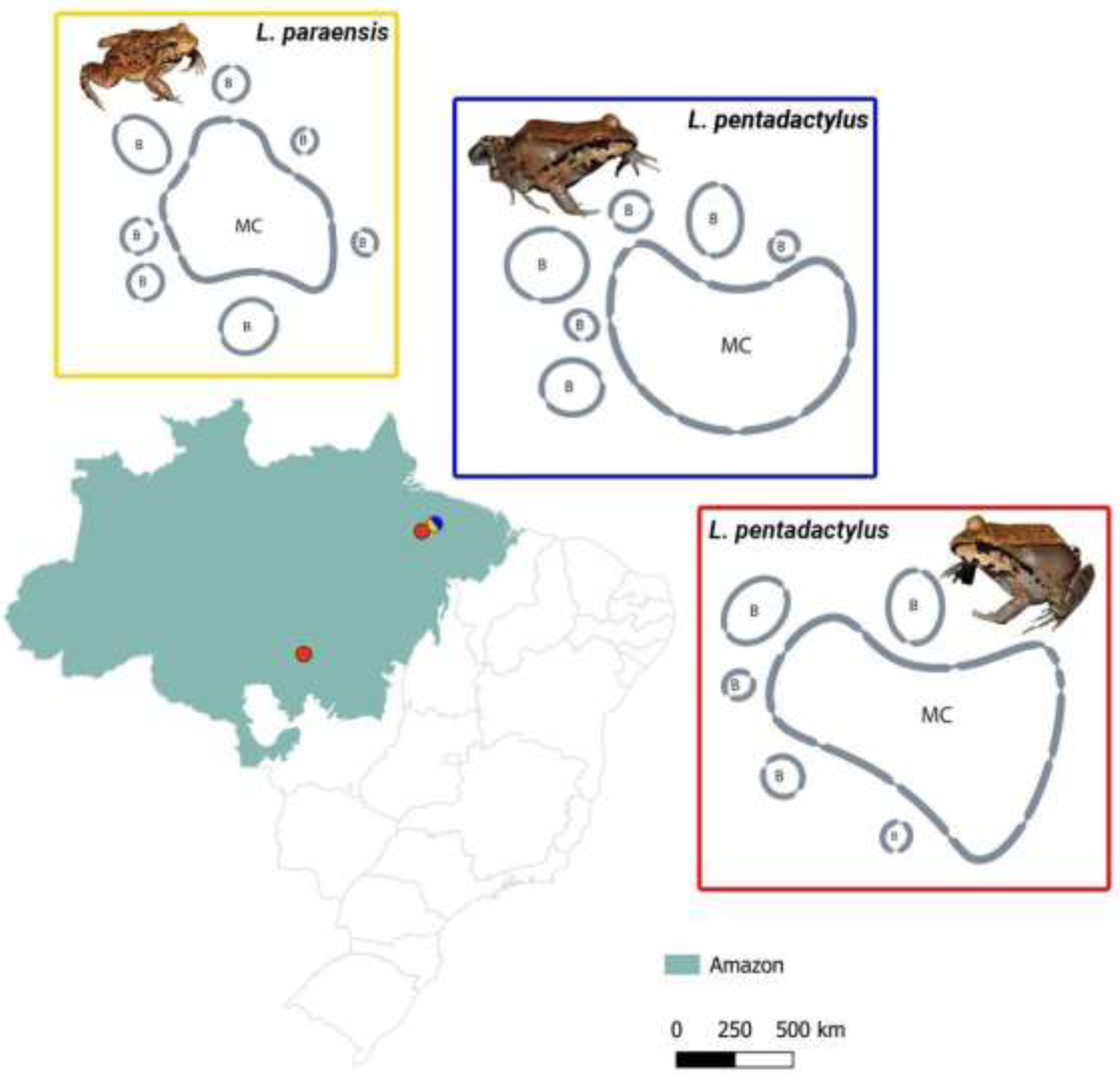
Map of Brazil highlighting the Amazon region and the sampling sites of the i) LPE populations so far analysed with a meiotic ring of 12 (red dots) or 10 (blue dots) chromosomes and ii) LPA with a ring of 8 chromosomes (yellow dots) in male meiosis. The other LPE population (red dots) with 12 sex chromosomes was previously analysed by ^15^ (2018) and ^17^ (2021), and the one with 10 sex chromosomes (blue) was previously analysed by ^21^ (2020). The chromosome identity numbers are indicated (B indicates bivalents and M chromosomes in the multivalents).

In the genus *Leptodactylus*, only the two species studied here have so far been shown to have systems with multiple sex chromosomes **(Supplementary Table 1)**, but most species remain unstudied, and more with rings may exist. If the absence of rings is confirmed by studies of other species, this would support the hypothesis that rings evolved more recently than the separation of LPE and LPA from other species. Independent evidence about the times when the different genome regions stopped recombining (such as Y-X divergence estimates for the different Y-X pairs) is, however, needed for definite conclusions concerning the age and rates of rearrangements. Below, we discuss the hypothesis that hybridization can help explain the occurrence of multiple rearrangements in a short evolutionary time.

Ring multivalents or chains must be initiated by a reciprocal translocation between two metacentric chromosomes, usually in centromere-proximal regions^2^, and further such events can add more chromosomes to the system. The events described here in *Leptodactylus* frogs are an example of the situation when the sex chromosome pairs are involved, as in the Monotreme mammals^14,18,19^. These are not sex chromosome turnovers, as the ancestral sex chromosome arms remain as part of the system, and as far as is known, the sex-determining system is not changed or moved to a different genomic location, as occurs in turnovers in other systems (reviewed in ^40^). We next discuss what factors may have allowed these neo-sex chromosomes to evolve, and what consequences this may have had, and how these effects may help to understand how the rings evolved, and shed light on Y chromosome degeneration.

### Pre-conditions allowing translocations to establish if they occur

In the introduction, we defined three main categories of factors that may influence the probability of a reciprocal translocation occurring and eventually being established in a population. Our findings corroborate the observation by^15^ that the chromosomes of these frogs exhibit localization of crossovers predominantly at the chromosomal termini, facilitating such translocations. ^21^ also analysed meiotic cells of LPE males with a ring of 10, using immunofluorescence to examine segregation (they also inferred possible effects of the dodecavalent on fertility). Our results are consistent with their observations of asynaptic regions in the centres of the chromosomes in the ring during anaphase I of male meiosis. In addition, the adjacent centromeres in the multivalent appear to segregate in an alternating pattern in male meiosis, a category (ii) factor that facilitates the production of balanced gametes^34^. Our results support the conclusion that such chiasma localization, and alternating segregation, were already present in the ancestral species, and therefore that one reason that these species have translocation rings is simply that the deleterious effects of the translocations are minor.

Localized recombination in males is expected in the ancestor, as well as after fusions, as terminal pairing is a precondition for correct segregation after a translocation, as mentioned above ^2^; also reviewed by ^41^). In most Anuran species so far studied, most chromosome pairs form ring bivalents in male meiosis, as pairing (and thus probably crossing over) is localized to the terminal regions. Genetic maps have confirmed strong terminal crossover localization in male, but not in female, meiosis of >30 species with male heterogamety, and 4 with female heterogamety, including in the families Ranidae, Hylidae, and Pipidae (as reviewed in ^42,43^), which is not specific to the sex chromosome pair. Species with female heterogamety may differ in the same way between male and female meiosis, as observations of chiasmata in lampbrush chromosomes in female meiosis of the Kajika frog *Buergeria buergeri* detected multiple crossovers distributed across the lengths of three of the four autosomal bivalents studied and concentrated at one end of the fourth bivalent ^44^. The terminal pairing observed in *Leptodactylus* (with male heterogamety), and crossover localization not specific to the sex chromosome pair may therefore reflect an ancestral genome-wide pattern in male meiosis. Whether or not such crossover localisation also occurs in female meiosis can be tested if this can be studied in the future, as in *B. buergeri*, or female genetic maps can be estimated.

### Possible genomic associations with translocation breakpoints

Analyses in several species have suggested the possibility that satellite sequences cause rearrangements such as the translocations studied here ^45–49^, and some studies have detected satDNAs in fusion or fission breakpoint regions^47,50–52^. This hypothesis is plausible, as tandem repeats can induce ectopic recombination^26^. For example, in house mice and humans, distinct chromosomes containing the same satDNA family have experienced Robertsonian translocations^53^, though this is not evidence that the satDNAs are involved unexpectedly often. They may simply be abundant in repeat-rich regions that recombine rarely, such as pericentromeric regions, as outlined in the Introduction. It therefore remains unclear whether the satDNA sequences analysed here are associated with the evolution of the multiple rearrangements that created the multivalents. Our *in situ* mapping in LPE demonstrated that most LpeSatDNAs are found near the ends of chromosomes, whereas the reciprocal translocations probably involved rearrangements with breakpoints in the middle regions, such that adjacent centromeres segregate in an alternating manner. This restricts the candidates for involvement of satellites to the few that are found in the middle regions (e.g., LpeSat03-152 and LpeSat09-1589).

TEs also cause genomic rearrangements, including deletions, insertions and translocations^54,55^. Large recent expansions of TEs might have preceded the rearrangements. For instance, DNA transposon EnSpm elements appear to be responsible for repeated translocations of sex-determining regions in various *Takifugu* pufferfishes ^56^. TEs inducing translocation of SD genes also occurs in salmonid fishes ^57^ and strawberry plants ^58^. Although these cases are very different from the one in the frogs studied here, TEs could have been involved in the rearrangements that formed the rings, and this might be detectable in the genome sequences. As a first step towards asking whether higher TE abundance is found in *Leptodactylus* species (either promoting the rearrangements, or accumulating as a consequence of them), we compared genome-wide TE abundance between two species in the genus, only one of which has multiple sex chromosomes (note, however, that these species are not close relatives and diverged long before the rings appear to have evolved, which post-dates the split with a much more closely related species that also lacks multiple sex chromosomes). Approximately 20% of the TE types detected may be species-specific (though this is an overestimate, because the use of the *L. fuscus* genome sequence for our TE library will have led to a failure to identify diverged sequences in LPE, possibly accounting for its overall low TE amounts, as explained above, and other reasons are also possible). It is also possible that TEs are ‘missing’ from LPE due to low read coverage ^59^.

Our preliminary TE landscape analysis does not suggest recent expansion of TEs in the LPE genome as a whole. Specific TE types may have expanded, but such expansions occur in most species that have been studied, including in *L. fuscus*, which has no ring system. Future analyses of the chromosomal locations of recent TE expansions in LPE could allow tests of whether these are associated with the formation of ring sex chromosomes. It is currently unclear whether TE accumulation (or accumulation of any specific TE types) occurred disproportionately in the breakpoint regions, which have not yet been identified in the species studied here.

Deeper sequencing and analysis of a reliable LPE genome assembly can potentially answer these questions, along with manual curation of the unknown TEs to help identify whether the genomes of lineages with rings carry genuinely lineage-specific TEs. Even with phased genomic sequence data to identify translocation breakpoints in multiple such species with suitable outgroups, however, it will be difficult to determine whether any TE-rich regions were the cause of the rearrangements, rather than reflecting accumulation after them.

### Effects of translocations on evolution of the chromosomes involved

It is currently unclear whether the translocations created new fully sex-linked (completely non-recombining) regions. Assuming that the ancestral pericentromeric regions in which the breakpoints probably occurred were not already completely non-recombining, the translocations may have created new regions in which recombination was reduced from the ancestral rarely recombining situation. These regions should then start accumulating repeats, including both satellites and transposable elements. In species with heteromorphic sex chromosomes, repetitive DNA sequences are often highly enriched on, or even, in some cases, limited to, one or both sex chromosome(s) (e.g., ^60–65^, reflecting their predicted accumulation in the absence of recombination ^61,66^. However, although several LpeSatDNAs are more abundant in one sex than the other (**Supplementary Table 2; Supplementary** Figure 8; Figure 2), none are completely X- or Y-specific. Other species with multiple sex chromosomes also exhibit minor, or no, differential accumulation of SatDNA sequences and heterochromatin in these multivalents ^67–70^. Occurrence of satellite sequences on several different chromosomes will obscure any sex chromosome specificity, and abundance differences among sampled individuals will also obscure sexual dimorphism.

If accumulation does occur on newly Y-linked regions, they should increase in size, while their X-linked counterparts should not, or should do so to a smaller extent. The cytogenetic results described here indicate expansion of one member of each pair of the ring chromosomes only for chromosomes 3 and 5 in LPE. The observation that most chromosomes in the two ring-forming frog species studied here have not become evidently heterochromatic (Figure 1), suggesting the possibility that none of the chromosomes in the ring contains large new non-recombining regions. Alternatively, these rings may be too young for cytologically detectable expansions to have occurred in most of them (though repeat accumulation in Y-linked regions is expected to occur rapidly after recombination is suppressed; see ^61^). However, it seems unlikely that all the neo-sex chromosomes of these species are very young, since fixation of multiple rearrangements in the male genome of a species probably requires a considerable evolutionary time, even in a small population in which genetic drift allows rearrangements to increase in frequency^32^. The lack of a large build-up of heterochromatin and repetitive DNAs might, however, be explained by natural selection against changes causing size differences that impede correct segregation in a ring system, increasing the frequency of deleterious unbalanced gametes ^2,71–73^. However, a build-up of repetitive sequences may have occurred, and could be detected by genome sequencing, even of it has not reached the level of detectability by staining for heterochromatin. Sequence data therefore have the potential to test whether recombination has completely stopped, or whether crossover events occasionally occur in the neo-sex chromosome arms.

If recombination has completely stopped, and sufficient time has passed since they did so, these regions should undergo genetic degeneration like that of other Y-linked regions, again making the loss of recombination potentially detectable. As noted in the Introduction, this has happened to all five Ys in the platypus and echidna, but it is currently unclear whether the frog neo-Y chromosomes have also done so. If only some have degenerated, it may be possible to determine which is the ancestral Y, and which are later additions to the system, and to estimate the time course of degeneration.

### Could large multivalents originate rapidly?

How old are these multivalents? We do not currently have data on sequence divergence between the species. The minimum age possible for the rearrangement that created shared elements of the LPE and LPA rings is therefore not currently known. We also do not have a sequence divergence between the different elements of the neo-Y and -X chromosomes (and, as already mentioned, cannot be certain that complete sex-linkage was created by the translocations); the ages when they evolved are not yet known. It is worth mentioning, however, that hybridization could potentially explain the occurrence of multiple rearrangements in relatively recent evolutionary history, as in the common shrew ^74^ and the black muntjac, *Muntiacus crinifrons* ^75,76^, although the exact mechanism of these multiple rearrangements is still unclear. If two populations’ chromosomes have undergone different translocations, their hybrids will be heterozygous for different rearrangements. Hybridization resulting in chromosomal polymorphisms, specifically inversions^77,78^ and frogs ^79,80^. Recurring Robertsonian translocations can produce substantial chromosomal chains in hybrid populations ^81–84^. Although hybridization has not been documented in *Leptodactylus*, it is plausible, as LPE and LPA are found in sympatry, and the conserved chromosome number in the genus would probably allow correct meiotic pairing and segregation in hybrids, and hybridization has been documented between other frog species ^86–89^. However, most elements of the ring are shared by both species, which suggests that this pre-date the split of the species.

## Material and Methods

### Samples, chromosomal preparation, DNA extraction, and C-banding

**Table 1** lists the species used in this study. *L. pentadactylus* and *L. paraensis* (hereafter referred to as LPE and LPA, respectively) were collected in sympatry in the Amazonian rainforest in northern Brazil, approximately 1,800 km from the site previously sampled by ^15^. Four additional species (designated *L. fuscus, L. mystacinus, L. latrans,* and *L. labyrinthicus*) were also examined, from distinct populations not previously examined. Individuals were identified based on species-specific morphological traits and deposited in the frog collection of the Federal Institute of Education, Science, and Technology of the State of Pará (IFPa) and in the Federal University of São Carlos (UFSCar). The specimens were collected from wild populations with authorization from the Chico Mendes Institute for Biodiversity Conservation (ICMBIO) and the System of Authorization and Information about Biodiversity (SISBIO) under license number 96067-1. The experiments followed the ethical standards set by the Federal University of São Carlos Ethics Committee on Animal Experimentation (CEUA) under process number 7994170423.

Mitotic chromosomes were obtained from bone marrow, following the protocol of ^85^, and meiotic chromosomes were obtained from males using the protocol of ^86^. Genomic DNA (gDNA) of LPE males and females was extracted using the phenol-chloroform method ^87^. C-positive heterochromatin was detected using the C-banding technique ^88^.

### Genome sequencing and satellitome analysis

The genomic DNAs of one LPE male and one female were sequenced on the BGISEQ-500 platform at BGI (BGI Shenzhen Corporation, Shenzhen, China) (PE-150) and yielded total sizes of 2.46 Gb and 2.43 Gb, respectively, which may represents an ∼1x coverage. The raw reads are available on the Sequence Read Archive (SRA-NCBI) under accession numbers SRR30896012 (male) and SRR30896011 (female). Satellite DNA (satDNA) sequences characterized from LPE individuals (see below) are deposited in the GenBank database under accession numbers (PQ462747 - PQ462850).

The satDNAs were identified following the pipeline described in ^89^ together with analysis with tools in the Tandem Repeat Analysis (TAREAN) package ^90,91^ available at https://galaxy-elixir.cerit-sc.cz/. First, low-quality reads were filtered using Trimmomatic ^92^, with the following parameters: LEADING:3 TRAILING:3 SLIDINGWINDOW:4:20 MINLEN: 100 CROP: 101. Then, 2×500,000 reads were randomly selected using SeqTK (https://github.com/lh3/seqtk) and analysed with TAREAN. Putative satDNA sequences identified in TAREAN were filtered from the raw reads using the DeconSeq tool ^93^, and this process was repeated until no new putative satDNA sequences were found. To avoid multigene family sequences identified by TAREAN, alignments with the putative satDNAs were conducted in GENIOUS 6.1.8 (Biomatters) and any sequences with similarity to multigene families were removed. To prevent redundancy within the set of satDNAs, we used a crossmatch search in RepeatMasker ^94^ and aligned each satDNA against the entire set of satDNAs. Genomic abundances were estimated using a RepeatMasker script (https://github.com/fjruizruano/satminer/blob/master/repeat_masker_run_big.py) by randomly selecting 10,000,000 raw reads and aligning them to the set of satDNAs. Divergence values for each member of each satDNA family from the family’s inferred consensus sequence were calculated for all site types using the calcDivergenceFromAlign.pl script ^89^ and the proportions of sequences with different abundances were plotted against the divergence values using the Kimura 2-Parameter correction for saturation. The satDNA families were named in order of decreasing abundance in the genome, following ^89^.

The genomes of neither ring-forming *Leptodactylus* species have been assembled, but a chromosome-level genome assembly is available for a closely related species *L. fuscus*. Genome and chromosome-wide synteny is very well conserved within frogs even across highly diverged (∼250MYA) groups, so we used the closely related species to infer the chromosomal distribution for the repeats ^39^. As mentioned above, this species is estimated to have diverged from LPE for about 27 MYA, and its karyotype is 2n=22, without meiotic rings; this assembly (NCBI accession GCA_031893025.1) was used to map LpeSatDNAs and discover their distribution on chromosomes.

### Polymerase Chain Reaction amplification of LpeSatDNAs

We designed primers for 18 LpeSatDNAs for *in situ* mapping and amplified them using polymerase chain reaction (PCR). From the 10 most abundant LpeSatDNAs, we excluded LpeSat007-21 due to its short motif length (21 bp). We also included 9 other sequences that, whether they were abundant or not, showed male-to-female (M/F) or female-to-male (F/M) abundance ratios > 1.4 and differed significantly from 1 (STATs), suggesting sex-biased pattern in abundance (**Supplementary Table 2**). The PCR protocol included an initial denaturation step at 95°C for 5 minutes, followed by 32 cycles with 95°C for 40 seconds, 60°C for 40 seconds, 72°C for 50 seconds, and a final extension at 72°C for 5 minutes. We confirmed amplification of the LpeSatDNAs by electrophoresis on 2% agarose gels and measured the concentration of the PCR products using a NanoDrop Spectrophotometer (ThermoFisher Scientific, Branchburg, NJ, USA).

### Preliminary repetitive DNA analyses in L. pentadactylus (LPE) and L. fuscus (LFU)

Currently, the only species in the genus with a chromosome-level genome assembly is *L. fuscus* (2n=22), presenting a haploid genome size of 2.3Gb and not exhibiting a ring sex chromosome system (GCA_031893025.1). Thus, to evaluate the repetitive sequence content in LPE (with no genome yet assembled) and compare it with that of *L. fuscus*, we first ran RepeatModeler2 ^95^ on the *L*. *fuscus* genome (GCA_031893025.1) with the -LTRStruc option enabled to include repeats ascertained based on their structures as well as their sequences. The resulting *de novo* transposable element (TE) library included sequences ranging from 27 to 13,275 bp. To quantify the repetitive fraction of the *L. fuscus* genome and the transposable elements in trimmed short-read data from LPE **(Supplementary** Figure 2), this library was used as input for RepeatMasker. As a rough indicator of the expansion history of the TE types, we used “repeat Landscape” analysis. This estimates Kimura 2-parameter (K2P) divergence values for each transposable element (TE) from the inferred consensus sequence of its type. To do this, we used the calcDivergenceFromAlign.pl script in the RepeatMasker package ^94^. To more accurately assess TE abundance in LPE males and females, we also used DNApipeTE, which is optimized for the classification and annotation of repetitive DNAs in low-coverage data (<1×) without requiring a genome assembly ^96^.

### Probe preparation and fluorescence in situ hybridization (FISH)

The 5S and 18S rDNA probes were obtained via PCR, following ^97^ and ^98^, respectively. Telomeric sequences (TTAGGG)n were generated by PCR using the (TTAGG)^5^ and (CCTAA)^3^ primers described by ^99^ in the absence of a DNA template.

All the aforementioned repetitive DNAs (i.e., rDNAs, telomeric, and the 14 amplified LpeSatDNA sequences) were labelled using the nick translation kit from Jena Bioscience (Jena, Germany), incorporating the fluorophores Atto488-dUTP (green) or Atto550-dUTP (red) according to the manufacturer’s instructions. Five satDNAs with motifs smaller than 40 bp (named LpeSat001-35, LpeSat002-35, LpeSat080-22, LpeSat091-37, and LpeSat096-32) were indirectly labelled with Biotin and further detected using Streptavidin-FITC (Sigma) (**Supplementary Table 3**).

FISH experiments were performed under high-stringency conditions ^100^. Each hybridization mix was composed of 200 ng of labelled probe plus 50% formamide, 2×SSC, 10% SDS, 10% dextran sulphate, and Denhardt’s buffer at pH 7.0 in a total volume of 20µl. The slides were dehydrated in ethanol (70%, 85%, and 100%) before counterstaining chromosomes with DAPI mounted in Vectashield (Vector Laboratories, Burlingame, USA).

### Comparative Genomic Hybridization (CGH)

CGH experiments were conducted to examine genetic differentiation between males and females of LPE. The gDNAs of males and females were labelled with Atto550-dUTP (red) and Atto488-dUTP (green), respectively, using the nick-translation protocol (Jena Biosciences, Jena, Germany) and further hybridized against the male mitotic and meiotic chromosomes. To block the common repetitive sequences, we used unlabelled C0t-1 DNA (gDNA enriched in highly and moderately repetitive sequences) produced according to ^101^. The hybridization mix was composed of 10µg of unlabelled female-derived Cot-1 DNA and 500 ng of both labelled male and female gDNAs. After using ethanol-precipitation, the pellet was air-dried and well mixed with 20μL of hybridization buffer (Denhardt’s buffer, pH 7.0), composed of 50% formamide, 2% 2×SSC, 10% SDS, and 10% dextran sulfate. The CGH experiments followed the methodology detailed in ^102^.

## Acknowledgements/Funding

This research was funded by the São Paulo Research Foundation (FAPESP) grant 2023/06898–0 (J.A.D.V.) and 2023/00955–2 (M.B.C.), and Brazilian National Council for Scientific and Technological Development (CNPq), grant number 302928/2021–9 (M.B.C.). W.-J. Ma is supported by an ERC starting grant (FrogWY, 101039501) and a starting grant from Vrije Universiteit Brussel (OZR4049). This study was financed in part by the Coordenação de Aperfeiçoamento de Pessoal de Nível Superior, Brasil (CAPES), Finance Code 001. All authors certify that they have no affiliations with or involvement in any organization or entity with any financial interest or non-financial interest in the subject matter or materials discussed in this manuscript.

## Author contributions

JADV, DC, WJM, and MBC conceived and designed research. JADV, RU, AJBG, RU and ABS conducted experiments. JADV, DC, WJM, QZ, RU, AJBG, FPF, TL, and MBC analyzed the data. WJM, ABS, and RU contributed with new methods. JADV, DC, WJM, QZ, RU, AJBG, ABS, FPF, TL, and MBC wrote the paper.

