## Supplementary material for "Cytological and preliminary genomic analysis of two *Leptodactylus* frog species (Anura, Leptodactylidae) with recently evolved large meiotic rings of multiple X and Y sex chromosomes": Supplementary Files.pdf

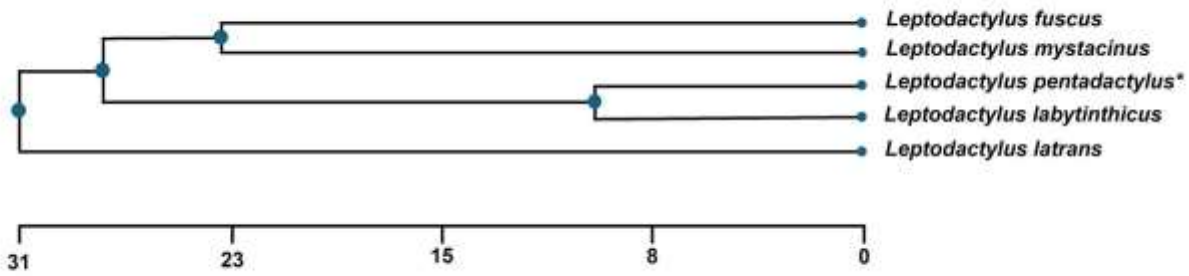


**Supplementary Figure 1.** Phylogenetic tree (http://www.timetree.og) highlighting the evolutionary relationships among selected *Leptodactylus* species. **indicates the only species with ring sex chromosomes*

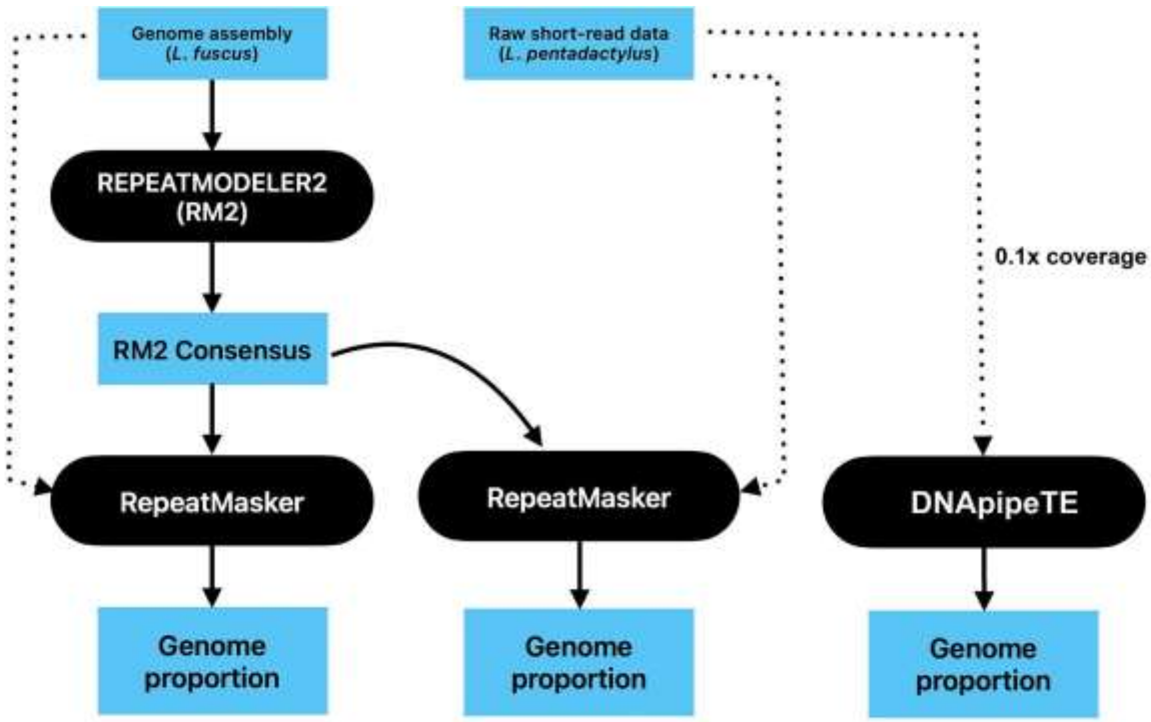


**Supplementary Figure 2.** Abstract of pipeline used to evaluate repetitive content from *Leptodactylus fuscus* and *Leptodactylus pentadactylus* species.

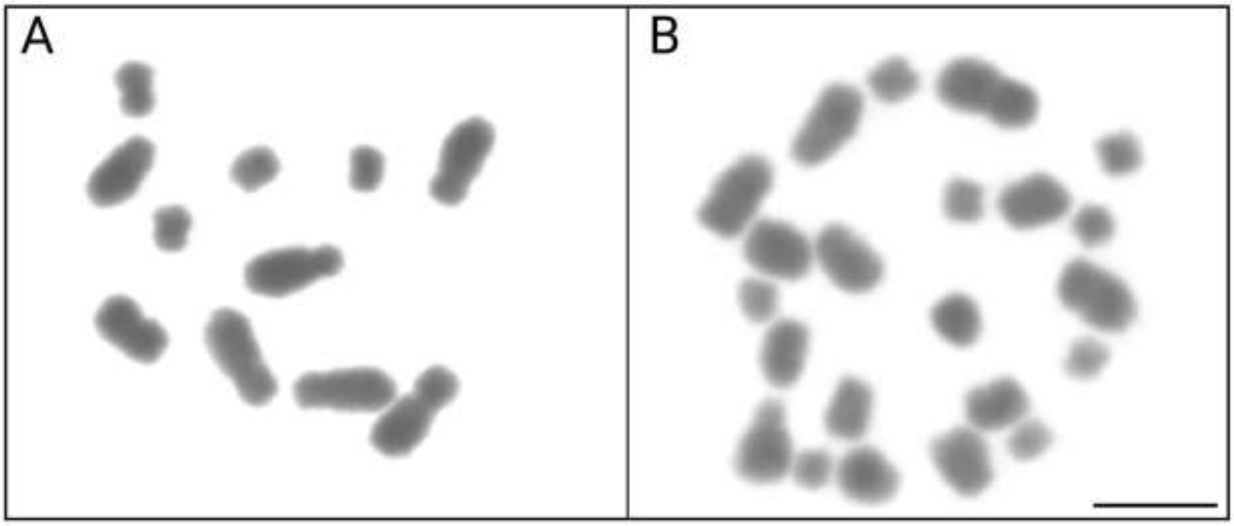


**Supplementary Figure 3.** Male meiotic cells from *Leptodactylus paraensis* (LPA). **A.** Metaphase meiotic II with 11 chromosomes. **B** Metaphase meiotic I with 22 chromosomes. Bar = 10 μm.

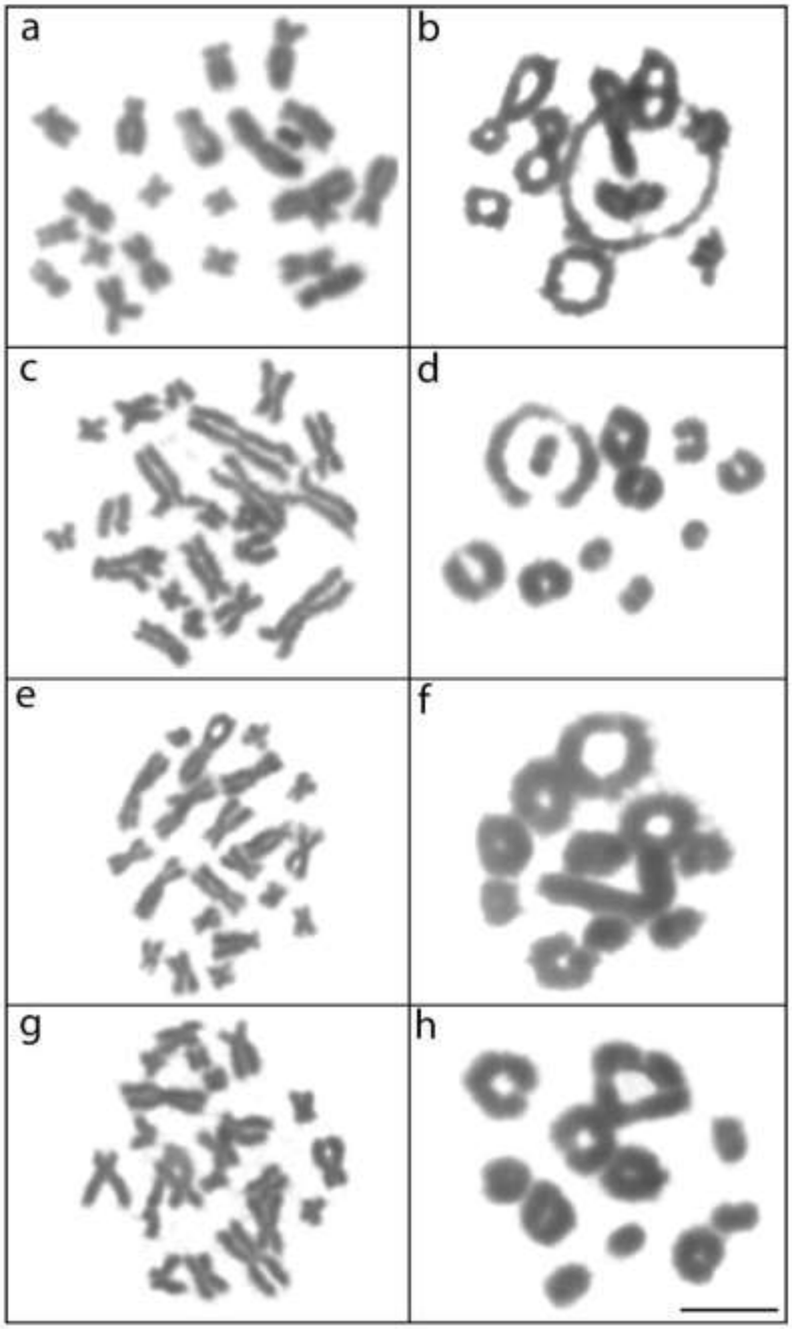


**Supplementary Figure 4:** Mitotic metaphases (a, c, e, and g) and meiotic metaphase I (b, d, f, and h) from *L. fuscus* (a and b), *L. mystacinus* (c and d), *L. latrans* (e and f), and *L. labyrinthicus* (g and h). Bar = 10 μm.

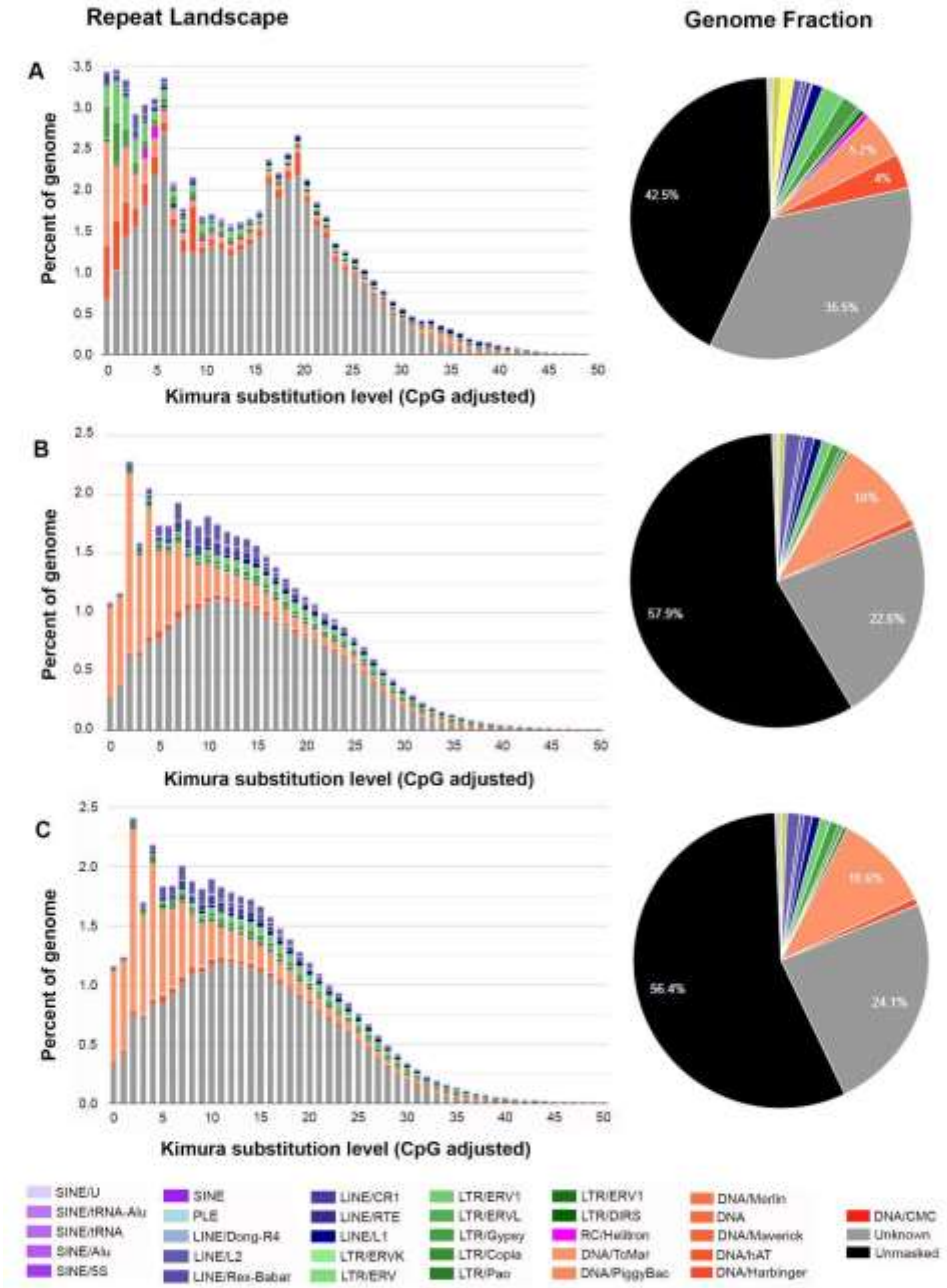


**Supplementary Figure 5**: Repeat landscape and pie chart of repetitive DNA class from *Leptodactylus fuscus* (A), *Leptodactylus pentadactylus* female (B) and male (C) using the RepeatModeler and RepeatMasker as described in the **Supplementary figure 2**.

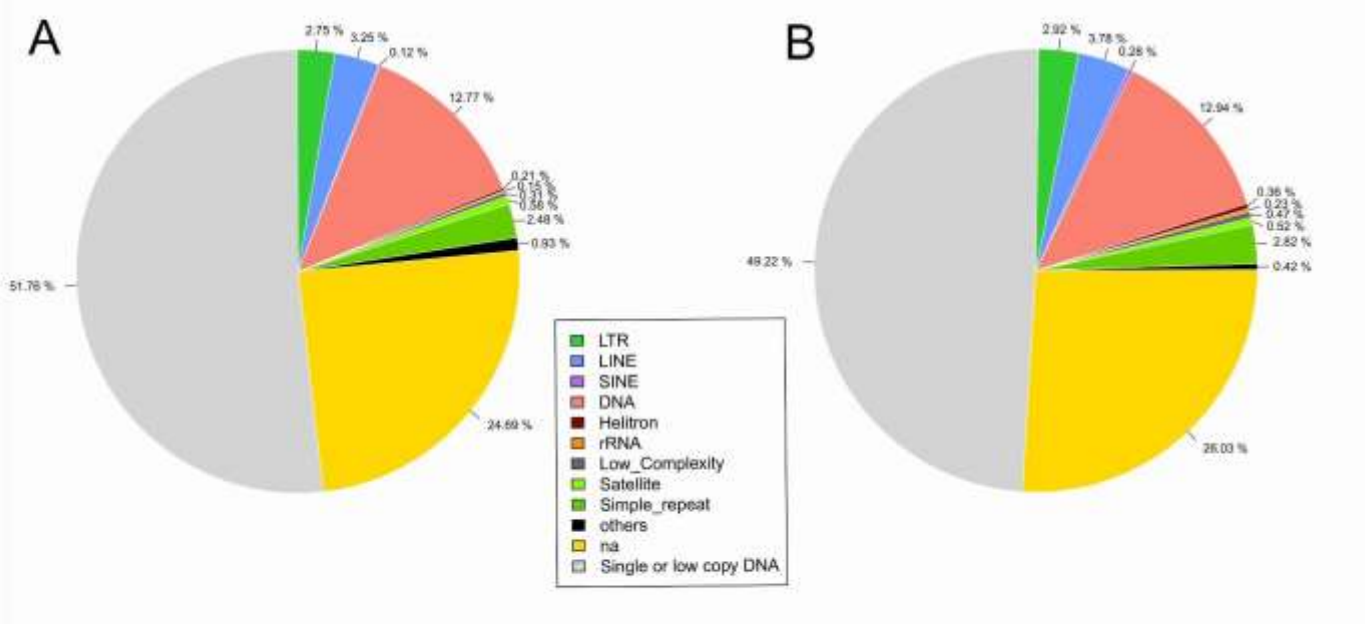


**Supplementary Figure 6:** Piechart showing the repetitive content in *Leptodactylus pentadactylus* male (A) and female (B) using the DNApipeTE described in **supplementary figure 2.** *Non-annotate (na).

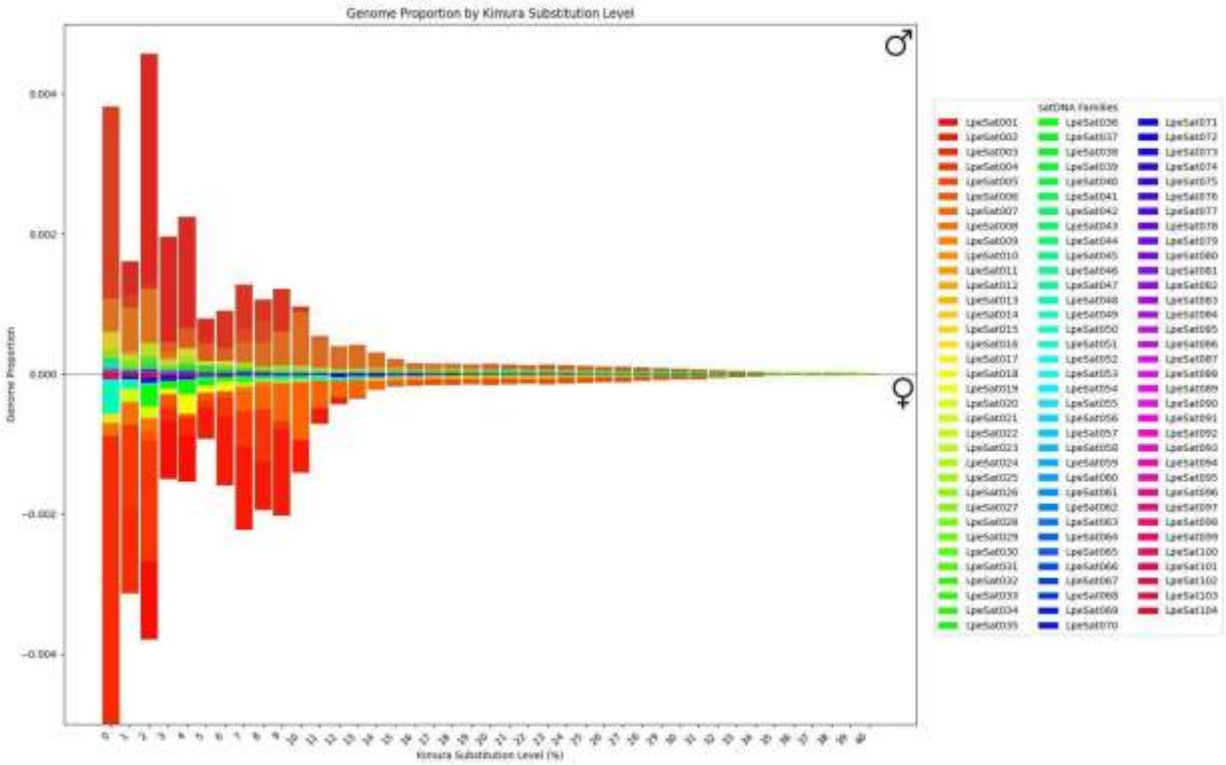


**Supplementary Figure 7:** Repeat landscape showing the genome abundance and divergence (Kimura substitution level) of LpeSatDNAs identified on male and female genomes.

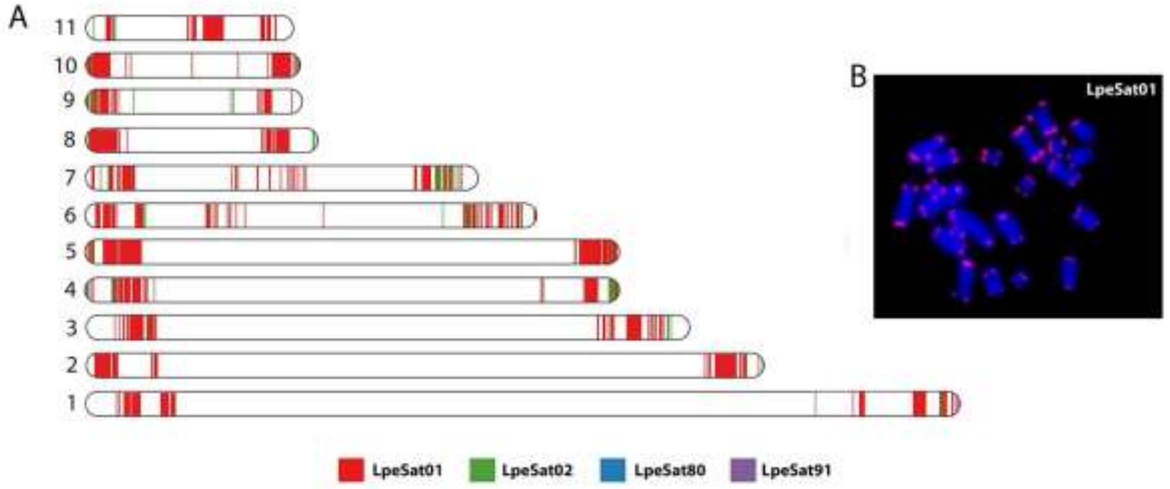


**Supplementary Figure 8:** A) In silico mapping of LpeSatDNAs to the genome of *L. fuscus* (GCA_031893025.1) underscoring the widespread and abundance occurrence of LpeSat01 in the terminal regions of all chromosomes. B) In situ mapping of LpeSat01 in a male metaphase plate of *L. fuscus.*

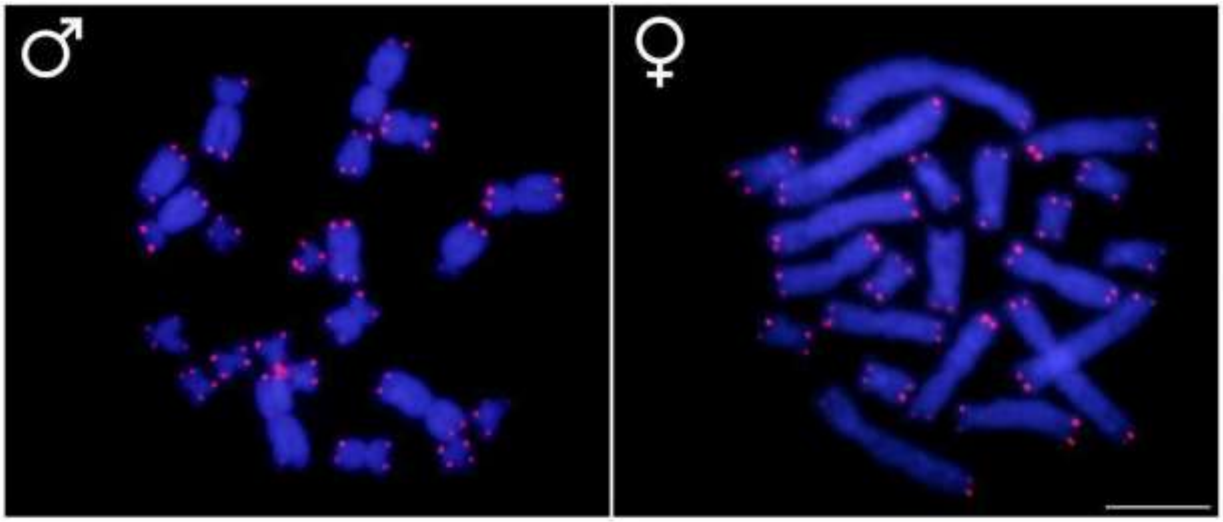


**Supplementary Figure 9.** Male and female mitotic cells of *Leptodactylus pentadactylus* hybridized with a telomeric probe. Bar = 10 μm.

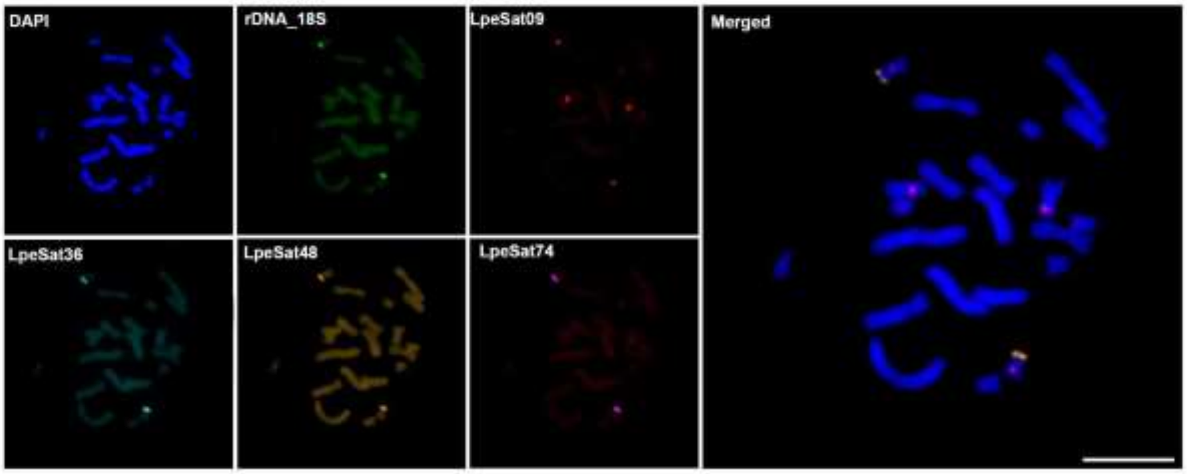


**Supplementary Figure 10.** Female mitotic cells from *Leptodactylus pentadactylus* sequentially hybridized with 18S rDNA 18S and several LpeSatDNAs, evidencing their co-localization on the same chromosome (pair 8). Bar = 10 µm.

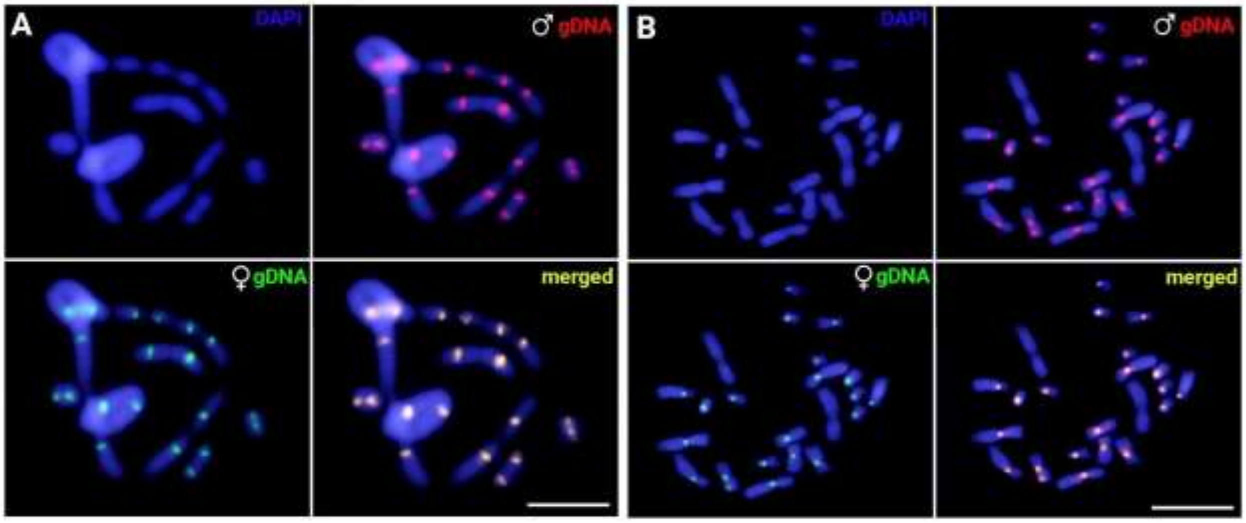


**Supplementary Figure 11.** Male meiotic (A) and mitotic (B) chromosomal preparations of *L. pentadactylus* after intraspecific comparative genomic hybridization (CGH). Chromosome spreads were probed with male- (red) and female-derived (green) genomic probes. The genomic regions with shared hybridization of both probes are yellowish. Scale bar = 10 µm.

**Supplementary Table 1.** Cytogenetic data for Leptodactylus genera available. The species with meiotic analysis are bolded. 2n = diploid number; NF = Nombre Fondamental; SCS = Sex chromosome system; MA = Meiotic analyses.

| Species | 2n | NF | SCS | MA | References |
|---|---|---|---|---|---|
| *Leptodactylus albilabris* | 22 | 44 | Ukn | NA | [103] |
| *Leptodactylus aff bokermanni* | 24 | Ukn | Ukn | **YES** | [104] |
| *Leptodactylus bolivianus* | 22 | 44 | Ukn | NA | [103] |
| *Leptodactylus bolivianus* | 22 | 44 | Ukn | NA | [103] |
| *Leptodactylus bufonius* | 22 | 44 | XY | NA | [105] |
| *Leptodactylus cf marmoratus* | **24** | 34 | Ukn | NA | [104] |
| *Leptodactylus chaquensis* | 22 | 44 | Ukn | NA | [15] |
| *Leptodactylus discodactylus* | 22 | NA | Ukn | NA | [106] |
| *Leptodactylus elenae* | 22 | 44 | Ukn | NA | [103] |
| *Leptodactylus furnarius* | 22 | 44 | Ukn | NA | [104] |
| ***Leptodactylus fuscus*** | **22** | **44** | **Ukn** | **YES** | [17], **present study** |
| *Leptodactylus gracilis delattini* | 22 | 44 | Ukn | NA | [107] |
| *Leptodactylus gracilis gracilis* | 22 | 44 | Ukn | NA | [107] |
| *Leptodactylus gracilis* | 22 | 44 | Ukn | NA | [103] |
| *Leptodactylus griseigularis* | 22 | 36 | Ukn | NA | [103] |
| *Leptodactylus hylaedactylus* | 26 | Ukn | Ukn | Ukn | [104] |
| ***Leptodactylus knudseni*** | 22 | 44 | Ukn | **YES** | [9] |
| ***Leptodactylus labyrinthicus*** | **22** | **44** | **Ukn** | **YES** | [34], **present study** |
| *Leptodactylus laticeps* | 22 | 44 | Ukn | NA | [103] |
| *Leptodactylus latinasus* | 22 | 42 | Ukn | NA | [103] |
| ***Leptodactylus latrans*** | **22** | **44** | **Ukn** | **YES** | [103], **present study** |
| *Leptodactylus leptodactyloides* | 22 | 36 | Ukn | NA | [103] |
| *Leptodactylus macrosternum* | 22 | 42 | Ukn | NA | [103] |
| ***Leptodactylus macrosternum*** | 22 | 44 | Ukn | **YES** | [17] |
| ***Leptodactylus marmoratus*** | **24** | 34 | Ukn | **YES** | [108] |
| *Leptodactylus melanonotus* | 22 | 44 | Ukn | NA | [109] |
| *Leptodactylus mystaceus* | 22 | 44 | Ukn | NA | [103] |
| ***Leptodactylus mystacinus*** | **22** | **44** | **Ukn** | **YES** | [38], **present study** |
| *Leptodactylus natalensis* | 22 | 38 | Ukn | NA | [103] |
| ***Leptodactylus ocellatus*** | 22 | Ukn | Ukn | **YES** | [110] |
| *Leptodactylus notoaktites* | 22 | 44 | Ukn | NA | [103] |
| *Leptodactylus paraensis* | 22 | 44 | **4X4Y** | **YES** | [103], **present study** |
| ***Leptodactylus pentadactylus*** | 22 | Ukn | **5X5Y** | **YES** | [10], **present study** |
| ***Leptodactylus pentadactylus*** | 22 | 44 | **6X6Y** | **YES** | [15] |
| *Leptodactylus petersii* | 22 | 42 | Ukn | NA | [103] |
| *Leptodactylus petersii* | 22 | 44 | Ukn | NA | [34] |
| *Leptodactylus plaumanni* | 22 | 44 | Ukn | NA | [107] |
| *Leptodactylus podicipinus* | 22 | 34 | Ukn | NA | [103] |
| ***Leptodactylus aff. podicipinus*** | **20** | Ukn | Ukn | **YES** | [34] |
| ***Leptodactylus podicipinus*** | 22 | 36 | Ukn | **YES** | [109] |
| *Leptodactylus pustulatus* | 22 | 44 | Ukn | NA | [103] |
| *Leptodactylus rhodomystax* | 22 | 44 | Ukn | NA | [34] |
| *Leptodactylus rhodonotus* | 22 | 44 | Ukn | NA | [103] |
| *Leptodactylus riveroi* | 22 | 42 | Ukn | NA | [103] |
| *Leptodactylus silvanimbus* | **24** | 48 | Ukn | NA | [103] |
| *Leptodactylus syphax* | 22 | 44 | Ukn | NA | [103] |
| *Leptodactylus troglodytes* | 22 | 44 | Ukn | NA | [103] |
| *Leptodactylus wagneri* | 22 | 36 | Ukn | NA | [109] |

**Supplementary table 2**. General features of *Leptodactylus pentadactylus* satellitome. The LpeSatDNA selected for FISH experiments are highlighted in bold.

| | | | Abundance | | | |
|---|---|---|---|---|---|---|
| **LpeSatDNA family** | **RUL** | **A+T%** | **Male** | **Female** | **M/F** | **Divergence** |
| **LpeSat01** | 35 | 71% | 0,014179 | 0,010638 | 1,332854353 | 3.87 |
| **LpeSat02** | 35 | 69% | 0,008699 | 0,013602 | 0,6395610634 | 9.55 |
| **LpeSat03** | 152 | 55% | 0,007955 | 0,02496 | 0,3187126625 | 2.08 |
| **LpeSat04** | 92 | 42% | 0,007118 | 0,012426 | 0,5727808764 | 6.79 |
| **LpeSat05** | 2216 | 51% | 0,005826 | 0,006415 | 0,9081191438 | 13.51 |
| **LpeSat06** | 2971 | 57% | 0,005164 | 0,005008 | 1,031035915 | 5.66 |
| LpeSat07 | 21 | 57% | 0,005132 | 0,004889 | 1,049613982 | 12.01 |
| **LpeSat08** | 97 | 58% | 0,004455 | 0,003079 | 1,44691215 | 2.98 |
| **LpeSat09** | 1589 | 58% | 0,003188 | 0,00391 | 0,8153829727 | 9.27 |
| **LpeSat10** | 5228 | 56% | 0,002457 | 0,002556 | 0,9611619277 | 11.02 |
| LpeSat11 | 4382 | 46% | 0,00228 | 0,001498 | 1,522164097 | 8.45 |
| LpeSat12 | 1205 | 60% | 0,002058 | 0,002235 | 0,9205658728 | 16.02 |
| LpeSat13 | 36 | 56% | 0,002024 | 0,001383 | 1,463472128 | 3.31 |
| LpeSat14 | 49 | 49% | 0,00191 | 0,001656 | 1,153429878 | 5.50 |
| LpeSat15 | 31 | 58% | 0,001763 | 0,001317 | 1,339149225 | 11.82 |
| LpeSat16 | 6930 | 56% | 0,00165 | 0,001636 | 1,00835934 | 5.35 |
| LpeSat17 | 577 | 57% | 0,00154 | 0,001733 | 0,8882055061 | 6.74 |
| LpeSat18 | 32 | 72% | 0,00145 | 0,002118 | 0,6844769905 | 5.63 |
| LpeSat19 | 44 | 59% | 0,001436 | 0,000774 | 1,855346071 | 4.26 |
| LpeSat20 | 31 | 71% | 0,001436 | 0,001457 | 0,9852910234 | 7.85 |
| LpeSat21 | 568 | 64% | 0,001335 | 0,001878 | 0,7107553132 | 2.70 |
| LpeSat22 | 4743 | 55% | 0,001318 | 0,001541 | 0,854832436 | 13.28 |
| LpeSat23 | 48 | 65% | 0,001251 | 0,001003 | 1,247060041 | 5.80 |
| LpeSat24 | 46 | 61% | 0,001248 | 0,00129 | 0,9676006196 | 8.43 |
| LpeSat25 | 72 | 60% | 0,000995 | 0,001011 | 0,9837593919 | 3.16 |
| LpeSat26 | 42 | 62% | 0,000905 | 0,00069 | 1,31153141 | 6.98 |
| LpeSat27 | 37 | 54% | 0,000814 | 0,000573 | 1,422371809 | 3.49 |
| LpeSat28 | 66 | 52% | 0,000806 | 0,001002 | 0,8048583727 | 4.17 |
| LpeSat29 | 37 | 62% | 0,000772 | 0,000765 | 1,009605763 | 3.23 |
| LpeSat30 | 39 | 56% | 0,000754 | 0,000734 | 1,026927731 | 6.75 |
| LpeSat31 | 84 | 64% | 0,000753 | 0,000558 | 1,349843164 | 18.62 |
| LpeSat32 | 39 | 56% | 0,00064 | 0,000362 | 1,768237995 | 7.17 |
| LpeSat33 | 1038 | 62% | 0,000625 | 0,000774 | 0,8067532999 | 22.09 |
| LpeSat34 | 31 | 61% | 0,00061 | 0,000282 | 2,16119492 | 9.10 |
| LpeSat35 | 50 | 70% | 0,000608 | 0,000495 | 1,226412222 | 7.89 |
| **LpeSat36** | 325 | 29% | 0,000587 | 0,001868 | 0,3142772489 | 4.87 |
| LpeSat37 | 72 | 49% | 0,000557 | 0,000722 | 0,7711051633 | 7.31 |
| LpeSat38 | 41 | 59% | 0,000551 | 0,000383 | 1,439617668 | 12.01 |
| LpeSat39 | 76 | 61% | 0,000547 | 0,000483 | 1,133230926 | 6.74 |
| LpeSat40 | 72 | 56% | 0,00054 | 0,000442 | 1,223167879 | 4.74 |
| LpeSat41 | 1677 | 55% | 0,000517 | 0,000523 | 0,9883594101 | 15.61 |
| LpeSat42 | 605 | 60% | 0,00045 | 0,000447 | 1,008096501 | 22.63 |
| LpeSat43 | 123 | 47% | 0,000433 | 0,000569 | 0,7617489005 | 3.16 |

| | | | | | | |
|---|---|---|---|---|---|---|
| LpeSat44 | 38 | 61% | 0,000395 | 0,000297 | 1,329992235 | 9.51 |
| LpeSat45 | 38 | 47% | 0,000392 | 0,000222 | 1,766613324 | 19.93 |
| **LpeSat46** | 108 | 33% | 0,000389 | 0,001256 | 0,3100862798 | 7.68 |
| LpeSat47 | 45 | 60% | 0,000379 | 0,000366 | 1,03409692 | 3.07 |
| **LpeSat48** | 178 | 33% | 0,000377 | 0,001352 | 0,2786608842 | 3.41 |
| LpeSat49 | 30 | 50% | 0,000377 | 0,000338 | 1,113195324 | 15.97 |
| LpeSat50 | 100 | 59% | 0,000338 | 0,000295 | 1,145452364 | 8.38 |
| LpeSat51 | 53 | 83% | 0,000335 | 0,000363 | 0,9235502703 | 12.79 |
| LpeSat52 | 30 | 57% | 0,00032 | 0,000257 | 1,243431614 | 12.91 |
| LpeSat53 | 383 | 63% | 0,00032 | 0,000238 | 1,343617226 | 7.07 |
| LpeSat54 | 99 | 54% | 0,000313 | 0,000278 | 1,124764553 | 6.81 |
| LpeSat55 | 31 | 52% | 0,000307 | 0,00025 | 1,229398627 | 10.47 |
| LpeSat56 | 45 | 67% | 0,000299 | 0,000356 | 0,8393590292 | 8.77 |
| LpeSat57 | 447 | 50% | 0,000293 | 0,000295 | 0,9934053866 | 12.01 |
| LpeSat58 | 30 | 43% | 0,000292 | 0,000177 | 1,648377147 | 5.57 |
| LpeSat59 | 37 | 59% | 0,000275 | 0,000418 | 0,6591347473 | 13.86 |
| LpeSat60 | 98 | 42% | 0,000266 | 0,000144 | 1,843022583 | 2.44 |
| LpeSat61 | 50 | 50% | 0,000255 | 0,000276 | 0,9245201567 | 4.44 |
| LpeSat62 | 42 | 50% | 0,000254 | 0,000255 | 0,9968065582 | 4.16 |
| LpeSat63 | 50 | 64% | 0,000251 | 0,000089 | 2,832278541 | 7.12 |
| LpeSat64 | 35 | 54% | 0,000242 | 0,000203 | 1,187861467 | 6.34 |
| LpeSat65 | 90 | 47% | 0,000232 | 0,000188 | 1,23473886 | 8.23 |
| LpeSat66 | 35 | 71% | 0,000227 | 0,000131 | 1,728533864 | 7.86 |
| LpeSat67 | 465 | 59% | 0,000217 | 0,000218 | 0,996588265 | 14.02 |
| LpeSat68 | 59 | 54% | 0,000216 | 0,000229 | 0,9400255048 | 8.77 |
| LpeSat69 | 38 | 50% | 0,000209 | 0,000198 | 1,05527617 | 8.16 |
| LpeSat70 | 30 | 50% | 0,000209 | 0,000286 | 0,7303831472 | 7.88 |
| LpeSat71 | 141 | 56% | 0,000204 | 0,000148 | 1,381412928 | 3.07 |
| **LpeSat72** | 70 | 53% | 0,000197 | 0,000097 | 2,035481152 | 9.45 |
| LpeSat73 | 46 | 59% | 0,000189 | 0,000142 | 1,333895907 | 5.73 |
| **LpeSat74** | 403 | 31% | 0,000179 | 0,000886 | 0,2016184388 | 7.04 |
| LpeSat75 | 67 | 61% | 0,000157 | 0,000075 | 2,089403001 | 2.42 |
| LpeSat76 | 112 | 65% | 0,000155 | 0,000113 | 1,379621441 | 4.42 |
| LpeSat77 | 41 | 46% | 0,000153 | 0,000183 | 0,8348121312 | 10.56 |
| LpeSat78 | 743 | 54% | 0,000146 | 0,000138 | 1,064089941 | 10.16 |
| LpeSat79 | 30 | 63% | 0,000143 | 0,000104 | 1,368710027 | 9.02 |
| **LpeSat80** | 22 | 50% | 0,000137 | 0,000055 | 2,480094189 | 9.68 |
| LpeSat81 | 35 | 66% | 0,00013 | 0,000124 | 1,048489964 | 6.05 |
| LpeSat82 | 830 | 56% | 0,000129 | 0,00009 | 1,436744307 | 4.24 |
| LpeSat83 | 40 | 68% | 0,000128 | 0,000075 | 1,695355365 | 9.16 |
| LpeSat84 | 27 | 59% | 0,000125 | 0,000077 | 1,63492693 | 7.61 |
| LpeSat85 | 38 | 55% | 0,000124 | 0,000097 | 1,287569341 | 9.23 |
| LpeSat86 | 46 | 54% | 0,000124 | 0,000124 | 0,9957757103 | 4.77 |
| LpeSat87 | 57 | 21% | 0,000117 | 0,000113 | 1,034625585 | 3.03 |
| LpeSat88 | 40 | 48% | 0,000116 | 0,000129 | 0,9012309023 | 5.36 |
| LpeSat89 | 31 | 77% | 0,000114 | 0,000139 | 0,8226507977 | 6.90 |
| LpeSat90 | 101 | 49% | 0,000112 | 0,000077 | 1,462881811 | 4.47 |
| **LpeSat91** | 37 | 59% | 0,000106 | 0,000045 | 2,35004712 | 11.83 |
| LpeSat92 | 217 | 65% | 0,000106 | 0,000123 | 0,8586067519 | 5.29 |

| | | | | | | |
|---|---|---|---|---|---|---|
| LpeSat93 | 585 | 62% | 0,000099 | 0,000155 | 0,6417960256 | 4.90 |
| LpeSat94 | 841 | 52% | 0,000098 | 0,000109 | 0,9013428882 | 2.01 |
| LpeSat95 | 45 | 62% | 0,000097 | 0,000071 | 1,367980281 | 9.87 |
| **LpeSat96** | 32 | 38% | 0,000095 | 0,000033 | 2,893865195 | 3.76 |
| LpeSat97 | 615 | 60% | 0,000084 | 0,000095 | 0,8856308975 | 4.05 |
| LpeSat98 | 26 | 62% | 0,000083 | 0,000092 | 0,8963277193 | 4.07 |
| LpeSat99 | 34 | 53% | 0,000083 | 0,000105 | 0,7885979334 | 8.73 |
| **LpeSat100** | 390 | 58% | 0,000078 | 0,000014 | 5,386677474 | 4.57 |
| **LpeSat101** | 74 | 54% | 0,000077 | 0,000011 | 7,327873404 | 4.40 |
| LpeSat102 | 216 | 45% | 0,00007 | 0,00008 | 0,8805428921 | 6.22 |
| LpeSat103 | 56 | 57% | 0,000069 | 0,000122 | 0,5658512868 | 3.29 |
| LpeSat104 | 208 | 53% | 0,000048 | 0,000042 | 1,151284662 | 11.27 |

**Supplementary Table 3**. List of primers used for the amplification of selected LpeSatDNAs.

| LpeSatDNAs | Forward primer | Reverse primer |
|---|---|---|
| LpeSat03-152 | TGAACTACGCGAAATGTGAT | AACTTCTCCGACTAGGGT |
| LpeSat04-92 | GTCACTGGGCACTAAACG | ATTAACGTGGCAATAACGCA |
| LpeSat05-2216 | ACTCAGGCAACCAGGTAT | TGCACAACCACAGACACT |
| LpeSat06-2971 | CATATCGGCTGTTGTAGG | ACACTATGTATGAACTGTGA |
| LpeSat08-97 | TCAGGATCAGTACAGGATA | TTATACTCCAGAGCTGCG |
| LpeSat09-1589 | TATGTGTCTGCTTCTGAGC | GACGGTTGGCAGAAGAAAT |
| LpeSat10-5228 | AAGCACCTGTCAGATGAGA | TGAGGATAGCATGGATGTC |
| LpeSat36-325 | AGCCAGAGATGACCGTGA | TCTGGCTACGAGGCTACT |
| LpeSat46-108 | TGCAGCTTGGCTTCACCC | CATCAATCCCAGGAAAGGG |
| LpeSat48-178 | TTAGAGCTGCATTTGCCC | AGAGCAAGAGCGCAGAGA |
| LpeSat72-70 | TCCCTTTGCCCACTAGAA | TGGTCCTTTAGCCCACTA |
| LpeSat74-403 | AGAATGCCAGACCTCGCT | TGACATCCTGTGTCTTCTTT |
| LpeSat100-390 | ACGTCGACTCATTGGAAG | GGGAAGTAAGGTGTTGGT |
| LpeSat101-74 | TACTACACAGACAGGACCA | CGTGCGGACATATTGTGT |
